# SLAy-ing oversplitting errors in high-density electrophysiology spike sorting

**DOI:** 10.1101/2025.06.20.660590

**Authors:** Sai Koukuntla, Tate DeWeese, Alexandra Cheng, Robyn Mildren, Aamna Lawrence, Austin R. Graves, Kathleen E. Cullen, Jennifer Colonell, Timothy D. Harris, Adam S. Charles

**Affiliations:** Department of Biomedical Engineering, Johns Hopkins University School of Medicine; Kavli Neurodiscovery Institute, Johns Hopkins University; Department of Neuroscience, Johns Hopkins University; Janelia Research Campus, Howard Hughes Medical Institute; Center for Imaging Science, Johns Hopkins University; Department of Otolaryngology - Head and Neck Surgery, Johns Hopkins University School of Medicine; Department of Neuroscience, Johns Hopkins University School of Medicine

## Abstract

The growing channel count of silicon probes has substantially increased the number of neurons recorded in electrophysiology (ephys) experiments, rendering traditional manual spike sorting impractical. Instead, modern ephys recordings are processed with automated methods that use waveform template matching to isolate putative single neurons. While scalable, automated methods rely on assumptions that often fail to account for biophysical changes in action potential waveforms, leading to systematic oversplitting of individual neurons into multiple putative units. Consequently, manual curation of these errors, which is both time-consuming and lacking in reproducibility, remains necessary. To improve efficiency and reproducibility in the spike-sorting pipeline, we introduce the Spike-sorting Lapse Amelioration System (SLAy), an algorithm that automatically merges oversplit spike units. SLAy employs two novel metrics: (1) a waveform similarity metric that uses a neural network to obtain spatially informed, nonlinear waveform representations, and (2) a cross-correlogram significance metric based on the earth mover’s distance between the observed and null cross-correlograms. To improve reproducibility and remove the need for manual tuning, we also develop an automatic parameter setting procedure for SLAy that accounts for dataset-specific characteristics. On simulated oversplitting across a diverse set of animal models, brain regions, and probe geometries, SLAy substantially outperforms an existing merging algorithm, achieving high recall without merging extraneous units. On the original datasets without simulated oversplitting, SLAy recovers ∼ 95% of merges found by human curators and human curators agree with ∼ 90% of merges suggested by SLAy. SLAy leverages multithreading for computational efficiency, running in less than 10 minutes for all recordings we tested. SLAy is also compatible with SpikeInterface, making it a practical and flexible solution for large-scale ephys data analysis across acquisition systems and spike sorters.

## 1 Introduction

Modern neuroscience strives to understand how the concerted activity of neuronal populations guides higher-order phenomena, including sensory perception and behavior. High-density electrophysiological (ephys) probes such as Neuropixels [1, 2, 3] have revolutionized these efforts by enabling chronic recordings of hundreds of neurons with single-neuron precision, driving new approaches to modeling multi-region population coding and dynamics [4, 5, 6]. Because ephys probes can reach deep brain regions and record at sub-millisecond resolution, they are particularly well-suited for studying neural processes that involve rapid, coordinated activity across multiple cortical and subcortical brain regions. Comparable techniques such as two-photon calcium imaging lack this capability due to limited imaging depth and slow calcium indicator kinetics. Indeed, Neuropixels probes have advanced our understanding of a wide variety of neural processes, including perception [7, 8], decision making [9, 10], and navigation [11, 12]. Extracting these insights from ephys recordings, which capture the joint electrical activity of many neurons, relies on spike sorting: identifying the timing and identity of individual neurons’ action potentials from the raw voltage traces. Although ephys recordings have historically been spike sorted by hand [13], the increasing channel count of high-density electrophysiology datasets — especially those recorded using multiple probes — renders manual spike sorting impractical and necessitates automated solutions.

Automated methods assume a probe records a distinct stereotyped voltage waveform for each neuron, determined by the neuron’s intrinsic biophysical properties and location relative to the probe. Early semi-automated methods involve three steps [14] (Fig. 1a, left): (1) thresholding to detect spikes, (2) dimensionality reduction, often using principal component analysis (PCA), to represent spike waveforms in a low-dimensional feature space, and (3) manual or automatic clustering in this feature space to identify putative neurons [15, 16]. While faster than fully manual clustering, these methods struggle with temporally overlapping spikes and typically require some manual intervention.

**Figure 1:**
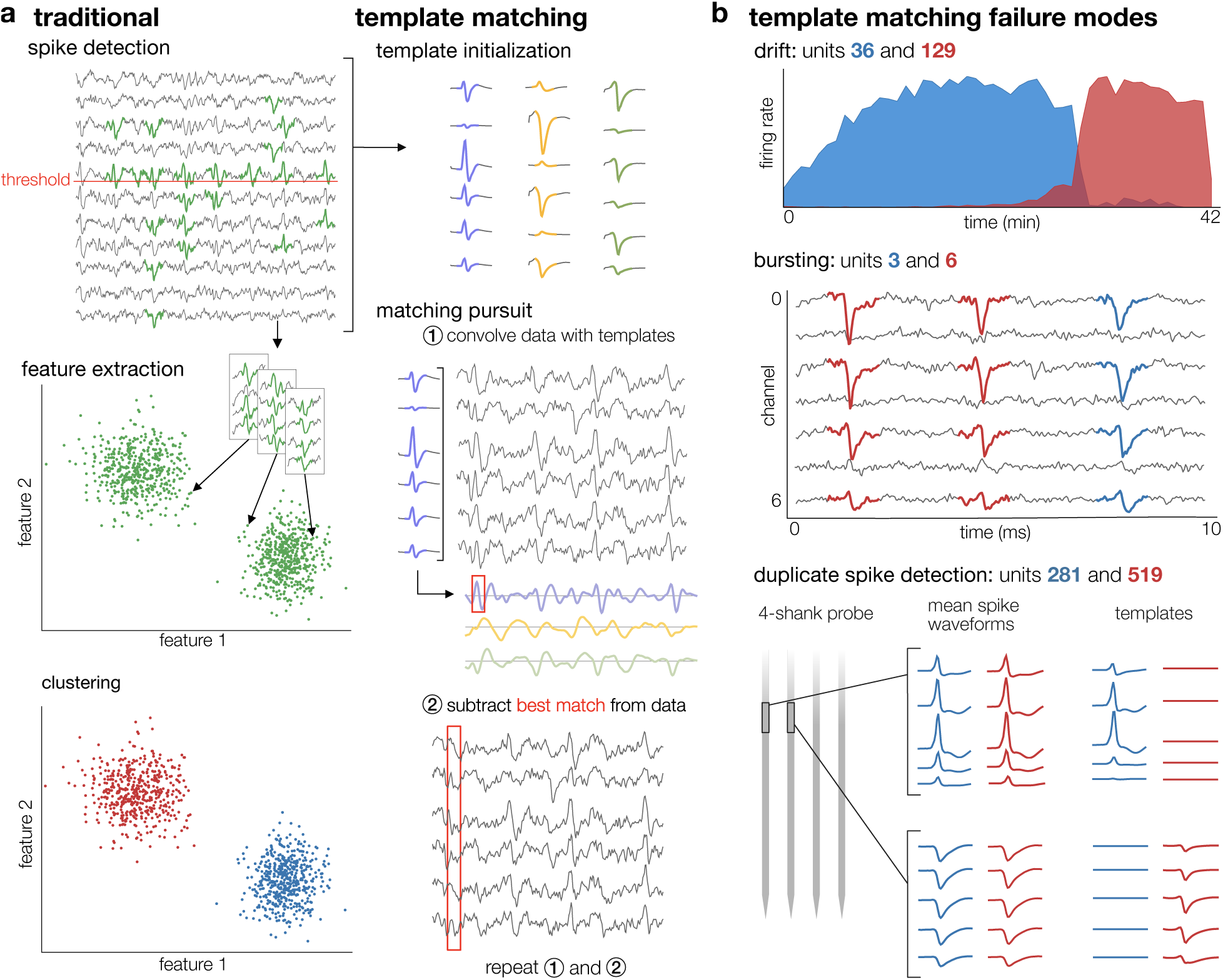
Spike sorting and errors in high-density electrophysiology data. **(a)** Left: Early automatic spike sorters involve three steps – spike detection, feature extraction, and clustering. Right: Newer template matching algorithms start with a template for each putative unit and alternate between detecting spike times with matching pursuit, and refining templates. **(b)** Common failure modes for template matching algorithms include waveform changes due to drift (top), biophysical waveform changes due to bursting (middle), and coincident spike time identification due to either improper template learning or detection of multiple parts of a single neuron (bottom).

Modern algorithms, including Kilosort [2, 17], SpyKING CIRCUS [18], and others [19, 20], employ template matching (Fig. 1a, right). They initialize a set of waveform templates for each putative neuron, then alternate between two steps: (1) detecting spikes by iteratively convolving the data with the templates and subtracting the best match (matching pursuit [21]), and (2) refining the templates based on the signal at the detected spike times. While template matching improves on earlier methods, it remains imperfect. Since template matchers assume a static waveform for each putative neuron (with rare exception [22]), they fail to account for waveform changes caused by probe movement or biophysical mechanisms (e.g. short-term facilitation during bursting; Fig. 1b), both of which are commonly observed in chronic recordings from behaving animals. As a result, template matchers often ‘oversplit’: they mistakenly fragment spikes from a (presumed) single neuron into multiple putative units.

Another class of oversplitting errors is the co-detection of a single spike by templates corresponding to multiple units, which results in many identical or slightly shifted spike times between units (Fig. 1b, bottom). This can occur for two reasons. In the more common case, the sorting algorithm learns an incomplete template that fails to capture the full amplitude or shape of a ground-truth neuron’s spike waveform. As a result, subtracting the template from the neuron’s spikes leaves behind a stereotyped residual waveform. The algorithm then learns another template for this residual, which is used to detect additional ‘spikes’ of a second unit at many of the original spike times (effectively ‘double-counting’ the same physical event). Alternatively, multiple compartments of the same neuron may be detected on noncontiguous parts of the probe. Because the templates used to detect spikes are constrained to be spatially localized, the sorter learns a separate template for each compartment. In both cases, the resulting units have highly similar mean waveforms (calculated by averaging the recorded voltage across the detected spike times) and a very large peak at the center of their cross-correlogram.

Overmerging errors, while also present, are much less common, since template matchers are intentionally biased towards oversplitting [17]. Although some template matchers include post-sort merging steps [17, 18], they rely heavily on waveform similarity to suggest merges and thus are unable to account for changes in waveform shape. Therefore, even state-of-the-art spike sorting algorithms require additional merging by a manual curator.

Manually correcting template matching errors primarily involves excluding low-quality units that correspond to multiple ground-truth neurons or noise, and merging oversplit units. Failing to address these errors can result in flawed scientific conclusions; analyses of single-neuron responses depend directly on the accurate assignment of spikes to neurons, and biased splitting may compromise the validity of dimensionality reduction and subsequent population-level analyses. Oversplitting in spike sorting has been shown to lead to underestimates of pairwise correlations between neurons [23], and similar errors in calcium transient attribution have been quantitatively shown to generate false correlations and misrepresent inferred coding properties [24, 25].

Although automated procedures for unit quality labeling have recently been developed [26, 27], no reliable automated method exists for merging oversplit units. The required manual curation is both time-consuming and lacking in reproducibility; curation decisions vary widely between curators due to factors including subjective judgment, fatigue, and differences in curation order [28, 16]. Growing interest in simultaneous multiprobe recordings [10, 29, 30] further exacerbates the limitations of manual curation, increasing both the time required and the potential for fatigue-related errors. Thus, automated methods for spike sorting error correction are essential for scalable and reproducible data processing pipelines.

To address the lack of a scalable and reproducible method for correcting template matching errors, we propose the Spike-sorting Lapse Amelioration System (SLAy). We first show that human curators consistently identify oversplitting errors in eight datasets that span a variety of brain regions, probe geometries, and animal models. We then introduce two key innovations in SLAy’s design: a waveform similarity metric that uses nonlinear neural-network-based representations of spike waveforms, and a cross-correlogram significance metric based on earth mover’s distance. SLAy also automatically tunes its parameters using realistic simulated oversplits, removing the need for manual parameter adjustment. On held-out simulated oversplits, we show that SLAy outperforms an existing merging algorithm, achieving near-perfect recall while avoiding extraneous merges. We also demonstrate that SLAy achieves high agreement with human curators—across datasets, human curators agree with ∼ 90% of merges suggested by SLAy, and SLAy independently identifies ∼ 95% of merges found by human curators. Critically, we show that traditional L2 waveform similarity and PCA features cannot achieve this level of performance across diverse datasets, highlighting the importance of neural-network-based waveform representations. By providing an automated method to correct template matching errors, SLAy stands to improve the efficiency of ephys data processing while maintaining the validity of downstream scientific conclusions.

## 2 Results

### 2.1 Oversplitting errors occur across brain regions, animal models, and probe geometries

To estimate the prevalence of oversplitting across recordings with a variety of characteristics, we sorted each of four chronic recordings with both Kilosort 2.5 (KS2.5) [2] and Kilosort 4 (KS4) [17], yielding eight datasets in total:

1. a macaque motor and premotor cortex recording from a *10 mm Neuropixels 1.0 probe (NP 1.0)* (data from [31]),
2. a macaque midbrain recording from a *prototype 128-channel Neuropixels NHP probe (NP NHP 128-channel)*,
3. a mouse hippocampus recording from a *Neuropixels 2.0 probe (NP 2.0)*, and
4. a rat insula recording from a *Neuropixels 2.0 probe*.

Each KS4 dataset was independently curated by two curators to quantify oversplitting and variability in manual curation. Though KS4 has largely superseded earlier Kilosort versions, we sought to explore whether oversplitting rates differed between Kilosort versions, specifically the previously widely used Kilosort version 2.5. Each KS2.5 dataset was thus curated by one curator. As is common practice in manual curation, curators considered only pairs of units with waveform similarity above a relatively low threshold. This reduces the number of candidate merges from all possible pairs of units to a more manageable subset.

Human curators identified oversplitting in seven out of eight datasets, with the fraction of units requiring merging ranging from 5–13% (Fig. 2a). The remaining dataset contained only 32 single-units. The two KS4 curators showed high (∼ 90%) agreement across datasets (Fig. 2b), indicating that the identified oversplits reflect genuine sorting errors rather than idiosyncratic judgments. The observed rates of oversplitting are nontrivial for downstream interpretation, as many analyses depend on accurate single-unit assignment and spurious splitting can obscure or, in the worst case, alter scientific findings [23]. Interestingly, no consistent pattern of oversplitting rates emerged between sorter versions; KS4 had higher, similar, or lower rates of oversplitting than KS2.5 depending on the recording. This suggests that the architectural changes in Kilosort 4 do not substantially reduce oversplitting errors.

**Figure 2:**
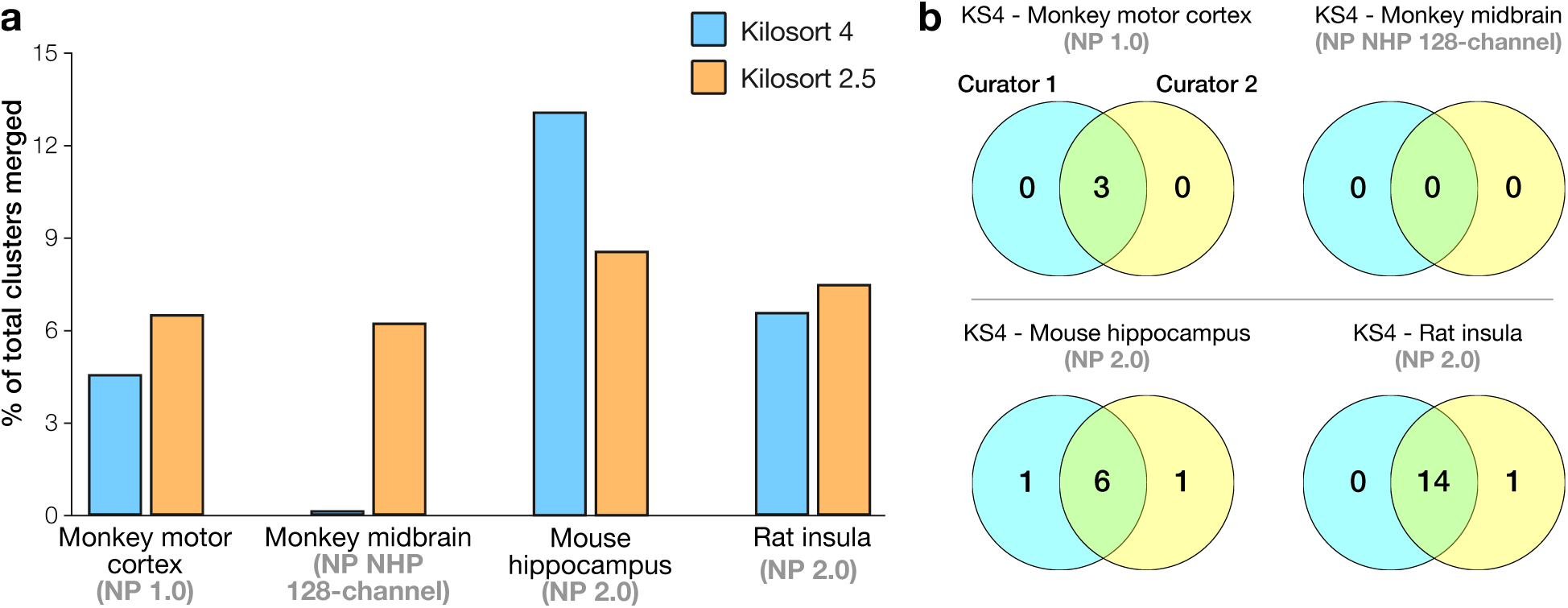
Oversplitting errors occur across brain regions, animal models, and probe geometries. **(a)** Fraction of total units merged by at least one human curator for KS4- and KS2.5-sorted datasets across four recordings with a wide range of characteristics. Curators found oversplits in 7 out of 8 of the datasets, indicating that oversplitting is widespread across datasets. **(b)** Venn diagrams showing the number of merges identified by each of two independent curators for KS4-sorted datasets. Human curators showed high agreement across datasets; only a few merges were identified by only one curator.

### 2.2 Quantifying waveform similarity, spike time dependence, and refractory period violations

Merging decisions, including those presented above, are currently made by human curators using graphical interfaces [32, 18] to qualitatively assess waveform similarity and cross-correlogram structure for each pair of units [33]. Our algorithm, SLAy, automates this time-consuming and subjective manual curation process through a combination of metrics that quantitatively capture the same criteria human curators use: waveform similarity, cross-correlogram structure, and refractory period violations.

#### 2.2.1 Waveform Similarity

A primary indicator of neural identity is the unit’s action potential waveform. When deciding whether to merge two units, curators consider one of two measures of waveform similarity. The first metric is the correlation, or, equivalently, the normalized inner product between *template* waveforms. Correlation provides a first-order measure of shape similarity and is used by spike sorting algorithms during post-sorting unit-merging steps [17], and by human curators to prioritize unit pairs to inspect manually. However, template similarity is only reliable when the learned templates accurately capture the spatiotemporal shape of spike waveforms, and popular spike sorters have nontrivial rates of inaccurate templates. Furthermore, Phy, the most widely used GUI for manual curation, uses templates to determine which channels to display for each unit [32]. Thus, improperly learned templates and the resulting waveform correlations can cause even diligent human curators to miss potential merges.

The second method involves projecting individual spikes from each unit into a low-dimensional feature space, typically through principal component analysis (PCA), and visually assessing the degree of overlap between the distributions for pairs of units. However, as principal components are computed independently for each channel, the resulting low-dimensional representations often fail to capture information about the spatial decay of waveforms. Combined with the tendency of human curators to consider only the principal components (PCs) for the one or two highest-amplitude channels, the per-channel feature approach can result in erroneous merging of two units with waveforms that are similar on their peak channels but decay differently across sites. Furthermore, since PCA representations span a low-dimensional linear subspace of the waveform space, they tend to smooth waveforms and remove clear distinguishing features that vary nonlinearly between units.

To address the shortcomings of these existing measures, SLAy computes waveform similarity in an expressive feature space obtained using an autoencoder (AE) (Fig. 3a). AEs can learn highly nonlinear features that represent the details of observed spike waveforms more faithfully than linear PCA approaches, thus preserving distinguishing features that can improve the accuracy of merge decisions. AEs consist of two neural networks: an encoder that generates a low-dimensional representation of its input and a decoder that reconstructs the input as accurately as possible from this low-dimensional representation. Because our AE learns a single representation for all channels rather than per-channel features as is common in PCA approaches, the AE features also capture the spatial decay of waveforms across channels.

**Figure 3:**
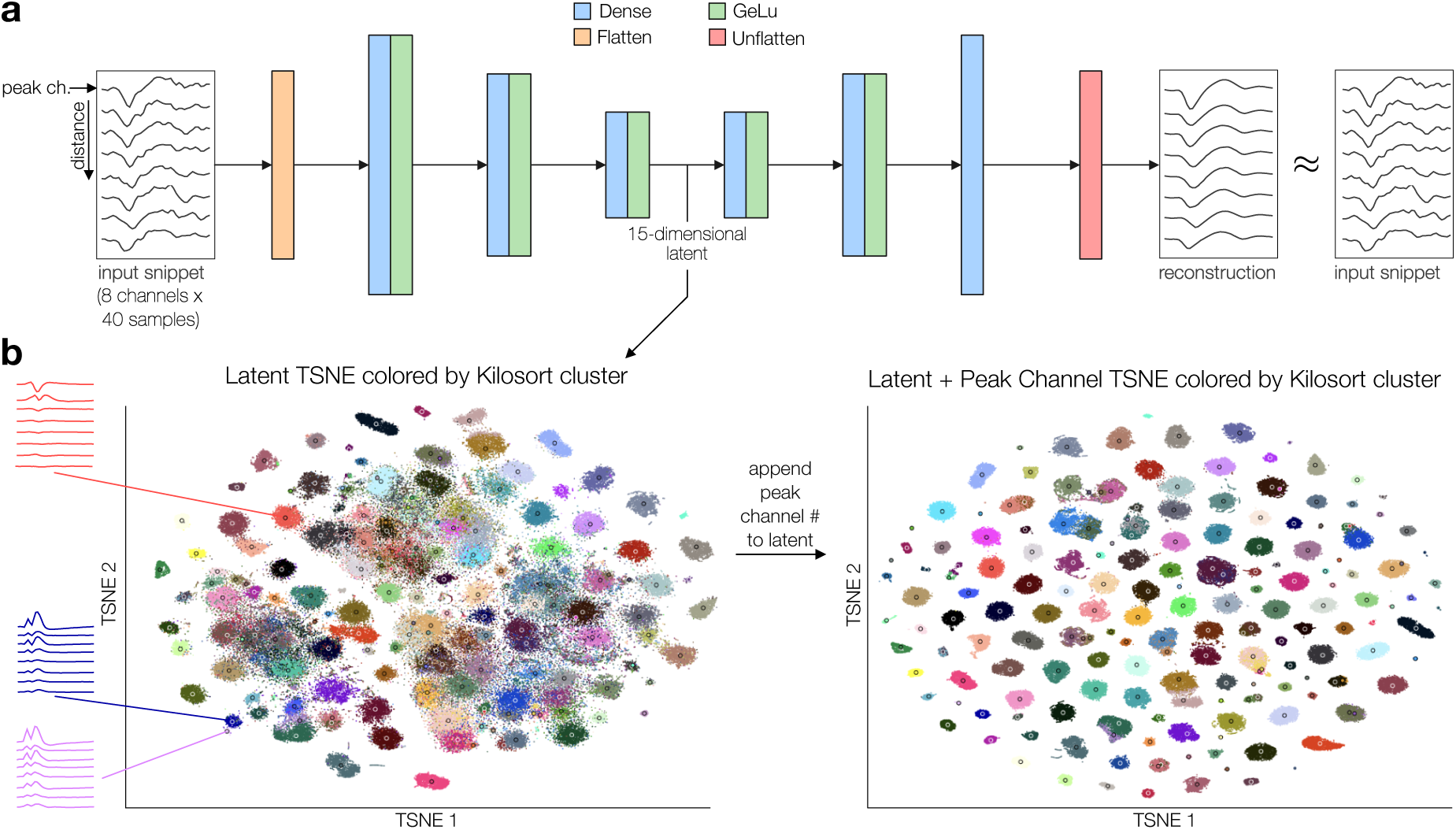
Autoencoder-based low-dimensional representations of spike waveforms. **(a)** We use an autoencoder network to obtain a low-dimensional nonlinear feature space in which to assess waveform similarity. **(b)** Left: The autoencoder latent space shows three large groups of units when using only waveform shape information, indicating relatively low waveform shape diversity. units close together in latent space have highly similar waveforms (blue and pink), and those far apart have different waveforms (blue and red). Right: Including information about the location of units on the probe improves unit isolation. units correlate roughly with Kilosort unit labels, though oversplitting is apparent in that some units contain data-points with multiple colors, representing multiple kilosort units.

To train our AE, we extract localized snippets of spike waveforms (8 channels × 40 samples) with the eight highest-amplitude channels sorted by proximity to the unit’s peak channel. Preprocessing the snippets in this way removes information about the spike’s absolute spatial position, ensuring that the autoencoder learns only features related to the waveform shape. Furthermore, limiting the snippet size rather than feeding in all probe channels reduces the number of network parameters, thus reducing the amount of data and computation time needed for training.

Once the AE is trained, we pass each spike waveform through the encoder to obtain their low-dimensional representations. Distances in the resulting latent space correspond to waveform similarity: spikes that are close together in the latent space have highly similar waveform shapes, while those far apart are markedly different (Fig. 3b, left). Because the autoencoder removes spatial information, the clusters in the latent space have considerable overlap, forming 3 main groups. If we reintroduce spatial information by appending the peak channel position to the low-dimensional latent vector (only for visualization purposes) results in much cleaner separation between units. This suggests some neurons recorded by the probe in different spatial positions have similar waveform shapes, perhaps due to having the same cell type and similar distance from the sites that capture them. Even in this 2D visualization of the AE latent space, oversplit or contaminated units are apparent (Fig. 3b, right). Furthermore, our AE achieves more accurate snippet reconstructions than PCA at the same dimensionality, highlighting the importance of nonlinear features for faithfully representing spike waveforms (Fig. S2).

To calculate the waveform similarity between two units from these features, we compute a similarity metric, *σ*, based on the Euclidean feature-space distance between the unit centroids relative to the standard deviations of the units’ spike latent vector distributions. Normalizing with respect to the spreads of each unit allows the metric to be robust to local distortions in latent space geometry — neural networks are known to warp latent space distances nonuniformly, such that equal distances in different regions need not reflect equal similarity. In practice, this normalization also reduces the sensitivity of the overall SLAy algorithm to changes in parameters (Fig. S1).

We use this normalized distance with an exponential kernel to constrain the metric to be between 0 and 1, with 0 corresponding to maximal dissimilarity and 1 corresponding to identical centroids. Because units with similar waveform shapes on different parts of the probe cannot correspond to the same ground-truth neuron, we also apply a multiplicative penalty based on the ratios of waveform amplitude at the two peak channels (Fig. 4, top left). The overall similarity metric is computed as

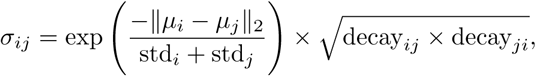

where *i, j* are the unit identities, *µ_i_* and *µ_j_* are unit centroids in the autoencoder latent space, 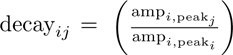, and decay 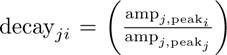. The use of a nonlinear encoder neural network means that the resulting waveform similarity metric accounts for spatial waveform decay and nonlinearly varying waveform features. Because the AE is trained directly on individual spike waveforms, our similarity metric avoids the shortcomings of computing similarity using templates, which may not faithfully represent waveform shape. Our similarity metric also takes into account the distribution of individual spike features for each unit, unlike simpler waveform similarity metrics based on the distances (e.g., L1 or L2 distance) between the mean waveforms. Importantly, the autoencoder similarity metric is not overly sensitive to randomness in the training process: correlations between unit similarities were very high for autoencoders trained with different random seeds (Tables S1, S2, see Methods Section 4.12), and SLAy suggested the same sets of merges across all random seeds tested.

**Figure 4:**
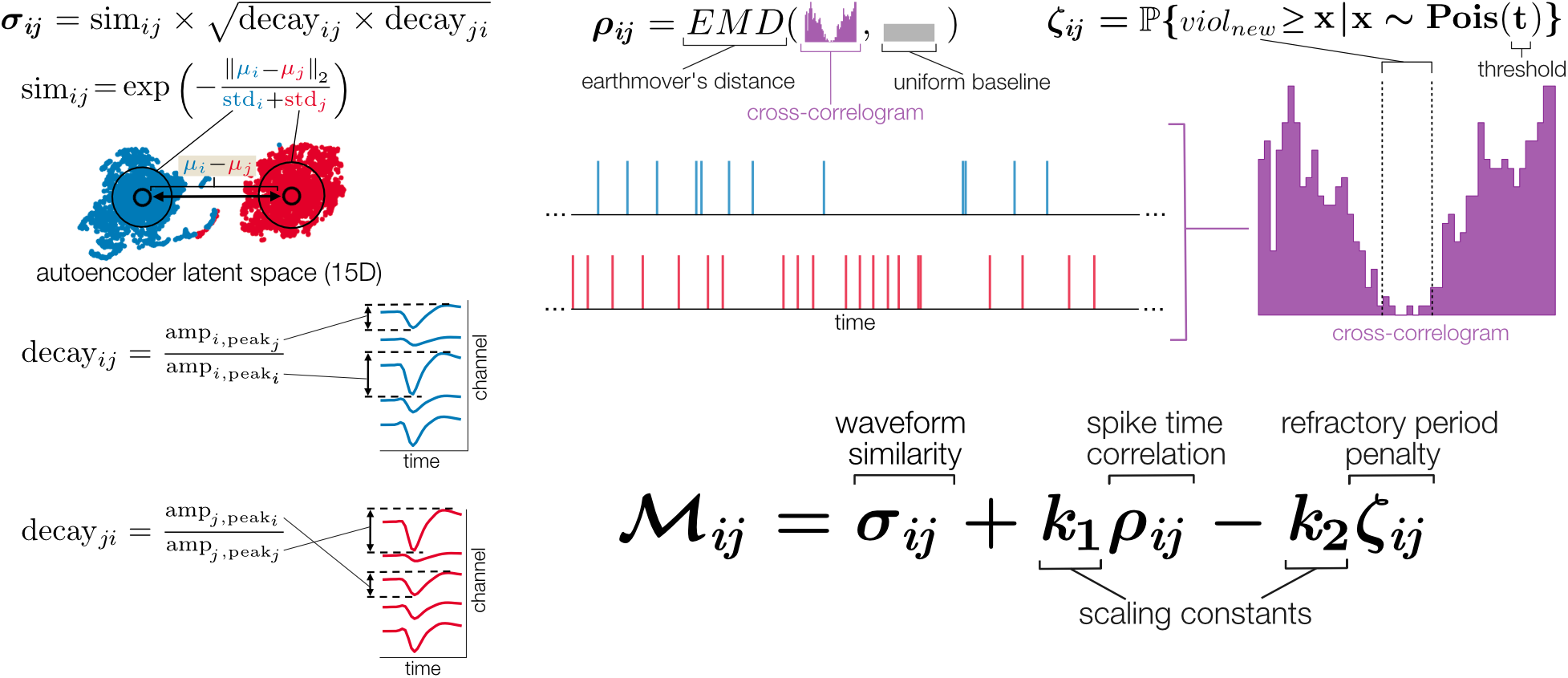
Metrics for automatic unit merging. We threshold a per unit-pair metric to determine which pairs to merge. The metric M*_ij_* between units *i* and *j* consists of three terms: waveform similarity (left), cross-correlogram structure (middle), and refractory period violation penalty (right). **Left:** unit similarity is defined as the spread-normalized distance between units in autoencoder latent space, scaled by the geometric mean of the waveform decay. **Middle:** Cross-correlogram structure is defined as the earth-mover’s distance (EMD) between the cross-correlogram and a uniform distribution. **Right:** Refractory period penalty is defined as the minimum contamination probability across a range of possible refractory periods. Bottom: parameters *k*_1_ and *k*_2_ control the relative influence of these measures on the M*_ij_*, and unit pairs with M*_ij_ >* a threshold are merged

#### 2.2.2 Cross-correlogram Shape Metrics

Although waveform shape is the primary criterion for merging units, curators must also consider the characteristics of the cross-correlogram (CCG). For example, spikes over the course of a burst can have substantial waveform variability due to the complex biophysics underlying action potential generation. Template matchers often assign these spikes to separate units, and waveform shape variability can make waveform similarity alone insufficient for merging these units. Therefore, merge decisions for bursting units must also account for significant peaks and dips in the CCG, which correspond to the bursting interspike interval (ISI) and refractory period, respectively. Conversely, a unit pair with very high waveform similarity is very unlikely to belong to the same ground-truth neuron if their CCG contains many spike collisions during the refractory period. To account for CCG characteristics when determining which units to merge, SLAy uses two metrics based on CCG shape: one that encourages merges between units whose CCG exhibits large peaks or troughs, and another that penalizes merges that would result in excessive refractory period violations.

No widely accepted method exists to determine whether a peak in a CCG is statistically significant, as human curators usually inspect CCGs visually to make qualitative judgments about their shapes. Some guides advise curators to merge units only if the CCG resembles the autocorrelograms (ACGs) of the individual units [33]. However, this approach can overlook potential merges, as structured (i.e., nonuniform) splitting due to bursting activity or drift-related changes in waveform shape can divide a single ground-truth neuron into units with distinct ACGs which also differ from the CCG.

One less prevalent approach is to perform binwise significance tests between the observed CCG and a “null” CCG, with a correction for multiple comparisons [34]. While this method does not assume that template matchers oversplit in an unbiased way, the statistical dependency between adjacent bins violates the independence assumption of many correction methods, which can result in underestimates of CCG significance. Additionally, bin-wise comparisons are highly sensitive to sampling noise and the choice of bin width.

To quantify the amount of structure in CCGs, we instead use the earth mover’s distance (EMD) between the (smoothed) observed CCG and a uniform distribution (Fig. 4, top middle). CCGs with pronounced peaks or troughs have large probability mass displacements relative to a uniform distribution and thus produce large EMDs. CCGs with random chance deviations from a uniform distribution instead yield low EMDs even if an individual bin passed a binwise significance test. However, this metric is only valid if the observed CCG resembles the “true” CCG given infinite observations. Because low-event CCGs are both unreliable estimates and far from uniform, we apply a multiplicative penalty to the metric for unit pairs that have CCGs with low spike counts. We also considered using the distance from the observed CCG to a per-unit-pair null CCG computed by repeatedly shuffling spike times. However, computing shuffled null CCGs was unnecessarily expensive, requiring many shuffling iterations while producing nearly identical metric values to a uniform distribution (Fig. S3).

To quantify the level of refractory period contamination, we use a modified version of the sliding refractory period (RP) violation metric, a standard metric for autocorrelogram refractory period contamination [27] (Methods). Given (1) the observed CCG, (2) a threshold of allowable contamination as a proportion of the baseline CCG event rate, and (3) a range of possible refractory periods, the sliding RP metric calculates the probability that the “true” CCG has an unacceptable level of contamination, assuming Poisson-distributed CCG events (Fig. 4, top right). We repeat this for each potential refractory period width, with the refractory period penalty defined as the minimum contamination probability across the set of refractory periods. Importantly, this metric is conservative when the baseline CCG even rate is low. For example, if the allowable refractory period contamination for a given CCG is 2 events and the observed CCG has 0 events in its refractory period, the sliding RP violation metric estimates that the CCG still has a ∼14% chance of being contaminated. This behavior is desired, since units should only be merged if there is statistical confidence that their CCG has a low rate of refractory period violations.

#### 2.2.3 Merge Decisions

To determine which unit pairs to merge, SLAy calculates a weighted sum of the three metrics described above: adding the waveform similarity (*σ_ij_*) and CCG significance metric (*ρ_ij_*), and subtracting the refractory period penalty (*ζ_ij_*) (Fig. 4, bottom).

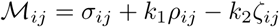

SLAy merges pairs of units with M*_ij_*above a threshold. The scaling constants *k*_1_*, k*_2_ ≤ 1 allow users to adjust the relative importance of each term. For example, in particularly bursty areas like hippocampus, a higher *k*_1_ could correct for burst-related oversplitting, while low-burst areas like cortex could have *k*_1_ set close to 0. *k*_2_ can be increased if unit isolation is especially important to downstream analysis. Though these parameters, including the merge threshold, can be tuned manually to align with a user’s judgment, in practice we recommend using the automatic parameter tuning scheme described below.

### 2.3 Realistic simulated oversplitting and automatic parameter selection

#### 2.3.1 Simulated oversplitting

We first sought to validate SLAy on oversplit recordings with known ground-truth. Such recordings could also serve as validation sets to tune the parameters of SLAy and other automatic merging algorithms. However, no such ground-truth datasets currently exist. Simulated high-density ephys recordings are not biologically realistic and spike sorters regularly achieve near-perfect performance on them [35]. Therefore, there are few errors to correct, and results on these datasets are not representative of real-world performance. Conversely, most real recordings do not have ground-truth for merging decisions, since manual curation is subjective. To address this issue, we developed a procedure to introduce simulated oversplitting errors into real spike-sorted recordings. This procedure performs one of four types of oversplitting on existing units in the sorting, based on the common classes of errors we have encountered: (1) amplitude-based, (2) burst-based, (3) drift-based, and (4) random (Fig. 5a; see Methods for details).

**Figure 5:**
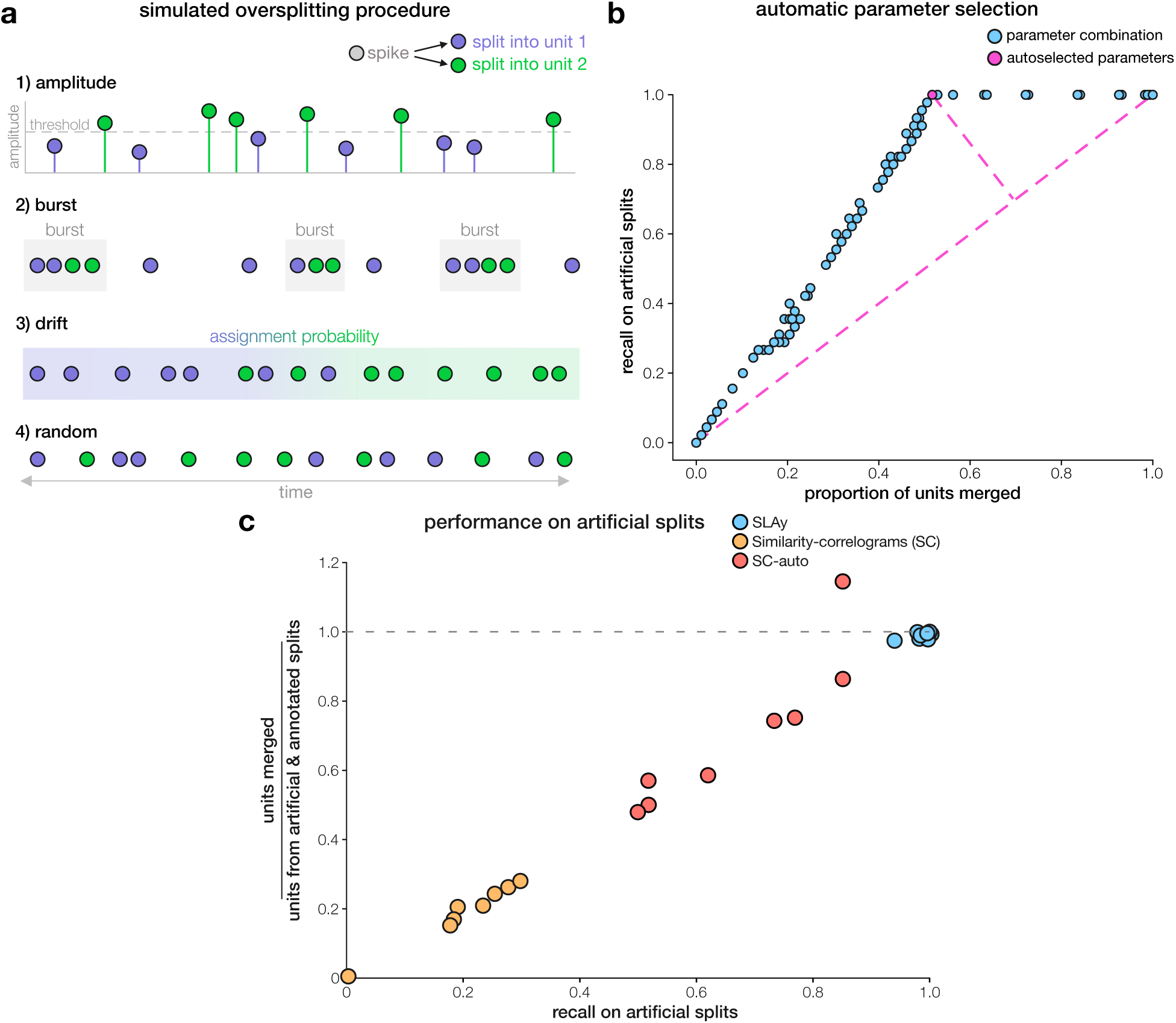
Simulated oversplitting, parameter selection, and evaluation. **(a)** We introduce four types of realistic oversplitting: (1) amplitude-based splits of a unit’s spikes based on a simple amplitude threshold, (2) burst-based splits of spikes in the second half of each burst into another unit, (3) drift-based splits of spikes based on their temporal position in the recording, and (4) uniform random assignment of spikes to new units. See Methods for full details. **(b)** To automatically select merging algorithm parameters for a specific dataset, we introduce simulated oversplitting and perform a parameter sweep. Our scheme selects the knee (pink) of the recall vs. proportion-of-units-merged curve, which is the parameter combination that achieves the highest recall while limiting the number of units merged. Pink dashed lines illustrate the knee-finding procedure: the knee is the point with the largest perpendicular distance to the line *y* = *x*. **(c)** Scatterplot of algorithms’ performance on artificial oversplitting. The y-axis shows the ratio between the number of units merged by the algorithm and the number of units from splits (artificial *and* real splits, with real estimated from human curation). A ratio *>* 1 indicates overmerging. Each data point corresponds to a single dataset-sorter combination and shows the method’s average performance across five draws of simulated oversplitting. Points near (1.0, 1.0) are jittered for visibility.

We split only a fraction of the total units in a given recording. Therefore, we can draw multiple ground-truth oversplit datasets from a single original dataset by varying the random seed of our splitting procedure. For example, a unit that is split on one draw may be unaltered on another, or a unit that undergoes an amplitude-based split on one draw may undergo a burst-based split on another. While all draws preserve the overall characteristics of the original recording (e.g., firing rates, waveform shapes, amplitudes), they are independent in terms of which units are split and the type of split applied to each. In practice, we found that a single draw of oversplitting was sufficient to automatically select parameters for SLAy – running SLAy with different combinations of autoencoder seed and simulated oversplitting draw resulted in autoselected parameters with mild variability (Tables 4, see Methods Section 4.12), but all runs produced the same set of suggested merges. We did, however, use multiple independent draws to benchmark different merging algorithms on each recording (see Section 2.4).

#### 2.3.2 Automatic parameter tuning

Because we have ground-truth information about the simulated splits, we can use them to evaluate the performance of automatic merging algorithms, including runs of the same algorithm with different parameters.

We leveraged this capability to develop an automatic parameter-tuning scheme, reducing the need for manual adjustment of SLAy’s parameters (Fig. 5a-b). On a given dataset, the scheme introduces simulated oversplitting and calculates the recall (fraction of ground-truth splits that were identified) and the proportion of total units merged for runs of the algorithm across many different parameter sets. Plotting these two performance metrics against each other produces a curve in the resulting 2D space (Fig. 5b, blue). Intuitively, a good merging algorithm should merge the oversplit units at a high rate while avoiding extraneous merges. We thus use a simple knee-finding procedure (greatest perpendicular distance to the line *y* = *x*) to identify the point on the curve (Fig. 5b, pink) where further relaxing the parameters would result in additional merges without improving recall on artificial splits.

Our oversplitting-based parameter selection scheme adds minimal additional runtime for SLAy. The similarity, cross-correlogram, and refractory period metrics are calculated independently of the coefficients and merge threshold. Thus, parameter sweeping involves only recombining these pre-calculated metrics with different coefficients, and then rethresholding. In practice, we found that a high refractory period penalty was necessary to avoid false-positive merges. Therefore, we fix *k*_2_ = 1 and use the automatic parameter tuning scheme to select the value of *k*_1_ and the threshold on the final metric. We do not tune parameters that affect the calculation of SLAy’s individual metrics, such as CCG bin size and refractory period width, as these are standardized by the field for each brain region.

The design of our parameter selection scheme confers two benefits. First, our simulated oversplitting procedure directly splits existing units, so the selected parameters will be well-tuned to the statistics (e.g., amplitudes, firing rates) of the specific recording of interest. Second, the scheme relies only on the recall and percent of units merged for each parameter combination, making it applicable to any merging algorithm. However, its effectiveness may depend on the design of the specific algorithm. Below, we show that our parameter selection scheme improves the recall of a prior merging algorithm at the cost of increased false positive merges.

### 2.4 SLAy achieves high recall and low extraneous merging on simulated oversplitting

Using the simulated oversplitting procedure described above, we generated datasets with ground-truth labels and evaluated the performance of the automatically tuned SLAy algorithm. For each combination of recording and Kilosort version analyzed in Section 2.1, we performed five draws of oversplitting and evaluated the resulting recall and the number of units merged. Because artificial oversplits were introduced into real recordings that already contained oversplit units, we compared the total number of units merged by the algorithm to the total number of split units. This total split count was computed as the sum of units involved in either artificial splits or curator-identified oversplits in the unaltered datasets. Importantly, the artificial oversplitting instances used for evaluation were generated using different random seeds than those used for automatic parameter selection. As established in the simulated oversplitting section, this ensures that the evaluation and tuning datasets share spiking characteristics while remaining independent, thereby avoiding train-test contamination.

SLAy reliably identified simulated oversplits across all recording–sorter combinations, achieving near-perfect recall across each of the eight datasets. This high recall was not the result of indiscriminate merging: the percentage of merges suggested by SLAy closely matched the percentage of total oversplits (Figs. 5c and S5, blue). The few simulated oversplits that SLAy failed to identify were usually rejected due to high refractory period penalties. In these cases, the cross-correlogram event count was low even in the non-refractory bins, making it difficult to confidently rule out refractory period violations. This reflects a deliberate design choice: the form of the refractory period penalty was selected to ensure that SLAy merges two units only when they are highly likely to have a low rate of refractory period collisions. Together, these results demonstrate that SLAy consistently corrects oversplitting errors while avoiding extraneous merges.

We also evaluated the performance similarity-correlograms algorithm (SC), an existing merging algorithm available through SpikeInterface. For each pair of units, the SC algorithm computes the mean waveform similarity and the similarities between the cross-correlogram (CCG) and each unit’s autocorrelogram (ACG), merging them only if all metrics exceed independent thresholds. As a result, this algorithm should in principle struggle to account for non-random oversplitting; in bursting neurons, for example, the first and last spikes in a burst can have considerably different waveforms, causing template matchers to systematically split spikes at the beginning and end of bursts into different units. A human curator can typically recognize an oversplit bursting neuron despite lower waveform similarity by noticing the large asymmetric peak in the CCG at the within-burst spike frequency. However, the SC algorithm would miss such an oversplitting error due to both low waveform similarity and low similarity between the asymmetric CCG and symmetric-by-definition ACGs. This limitation suggests that the SC algorithm would struggle to achieve high recall on our simulated oversplits.

This was indeed the case: the similarity-correlograms algorithm with default parameters was overly conservative across all recordings and Kilosort versions, performing far fewer merges than the number of simulated oversplits (Fig. 5c, yellow). As a result, it achieved a maximum recall of only ∼ 0.3 across datasets (Fig. 5c, yellow). To ensure that SLAy’s parameter selection scheme did not provide an unfair advantage, we also evaluated the performance of similarity-correlograms with our automatic parameter selection procedure (SC-auto). Our parameter selection scheme improved SC-auto’s recall on simulated splits across datasets (Figs. 5c and S5, red; *t*(7) = 7.50, *p <* 0.001*, d* = 2.65). For example, on the monkey motor cortex recording sorted with Kilosort 4, using autoselected parameters improved mean recall across draws from 0.17 to 0.9, about a fivefold increase. However, this improvement in recall was at the cost of extraneous merges – SC-auto merged more units than the total number of split units on one recording–sorter combination. We will also show later that on unaltered data, SC-auto makes extra merges that human curators disagree with. Even with recall-optimized parameters, SC-auto did not attain near-perfect recall for any dataset (Fig. 5c, red), with recall as low as ∼ 0.5 on the mouse hippocampus recording sorted with Kilosort 4 (Fig. S5). We suspect that the apparent ceiling on the SC-auto recall values for some datasets is due to an overly conservative contamination check in SpikeInterface’s automerging function, which rejects any merges that would increase the units’ contamination metrics even slightly.

SLAy achieved significantly higher mean recall across datasets than SC (*t*(7) = 24.19, *p* ≪ 0.001*, d* = 8.55) and SC-auto (*t*(7) = 5.51, *p <* 0.001*, d* = 1.95). Furthermore, SLAy achieved recall higher than (39/40) or equal to (1/40, perfect recall for both) SC-auto across every individual draw of simulated oversplitting for all recording–sorter combinations tested. Thus, SLAy consistently outperforms the primary existing automerging algorithm, even when that algorithm is augmented with our parameter tuning scheme.

### 2.5 SLAy achieves high agreement with human curators on unaltered real datasets

Having evaluated SLAy and an existing algorithm on hybrid datasets with ground-truth split information, we sought to evaluate how well SLAy and similarity-correlograms aligned with human curators. We applied SLAy (including autoselected parameters), similarity-correlograms (SC), and similarity-correlograms with autoselected parameters (SC-auto) to the original, unaltered sorted recordings. Additionally, to investigate whether SLAy’s autoencoder-based waveform similarity provides a practical advantage over traditional measures, we tested two ablated versions of SLAy: 1) a version that replaces the autoencoder similarity with the normalized L2 distance between mean waveforms (SLAy-L2), and 2) a version that replaced the autoencoder latent space with 15-dimensional PCA space (SLAy-PCA) but retained all other aspects of the similarity calculation, including the per-unit spread normalization. We applied automatic parameter selection to these version of SLAy, for which the cross-correlogram-related SLAy metrics remained unchanged.

For each algorithm, we computed the percent of human merges that the algorithm successfully recovered (recall) and the percent of the algorithm’s merges that a human curator agreed with (precision). For KS2.5 datasets, this was simply the percent of the algorithm’s merges found by the sole curator. For KS4 datasets, it was the percent of the algorithm’s merges found by *either* of the two curators. Some disagreements between human curators and algorithms could have stemmed from confounding factors, e.g., curator fatigue or limitations of the curation interface. To address this, a human curator re-evaluated any automated merges that they had not identified initially. Crucially, to avoid bias during this secondary review, we included additional unit pairs not suggested by any algorithm as foils; the curator was thus blinded to whether each merge was actually suggested by an algorithm.

SLAy’s identified merges aligned closely with those of human curators (Fig. 6a). Across datasets, SLAy found ∼ 97% of merges that were initially identified by at least one human curator, missing only one curator-identified merge (Figs. 6b, S6, S7). SLAy did not achieve this performance by simply identifying a large number of merges — after blinded re-evaluation of SLAy’s merges, both human curators agreed with ∼ 90% of all merges suggested by SLAy, and at least one human curator agreed with ∼ 95% of SLAy’s merges.

**Figure 6:**
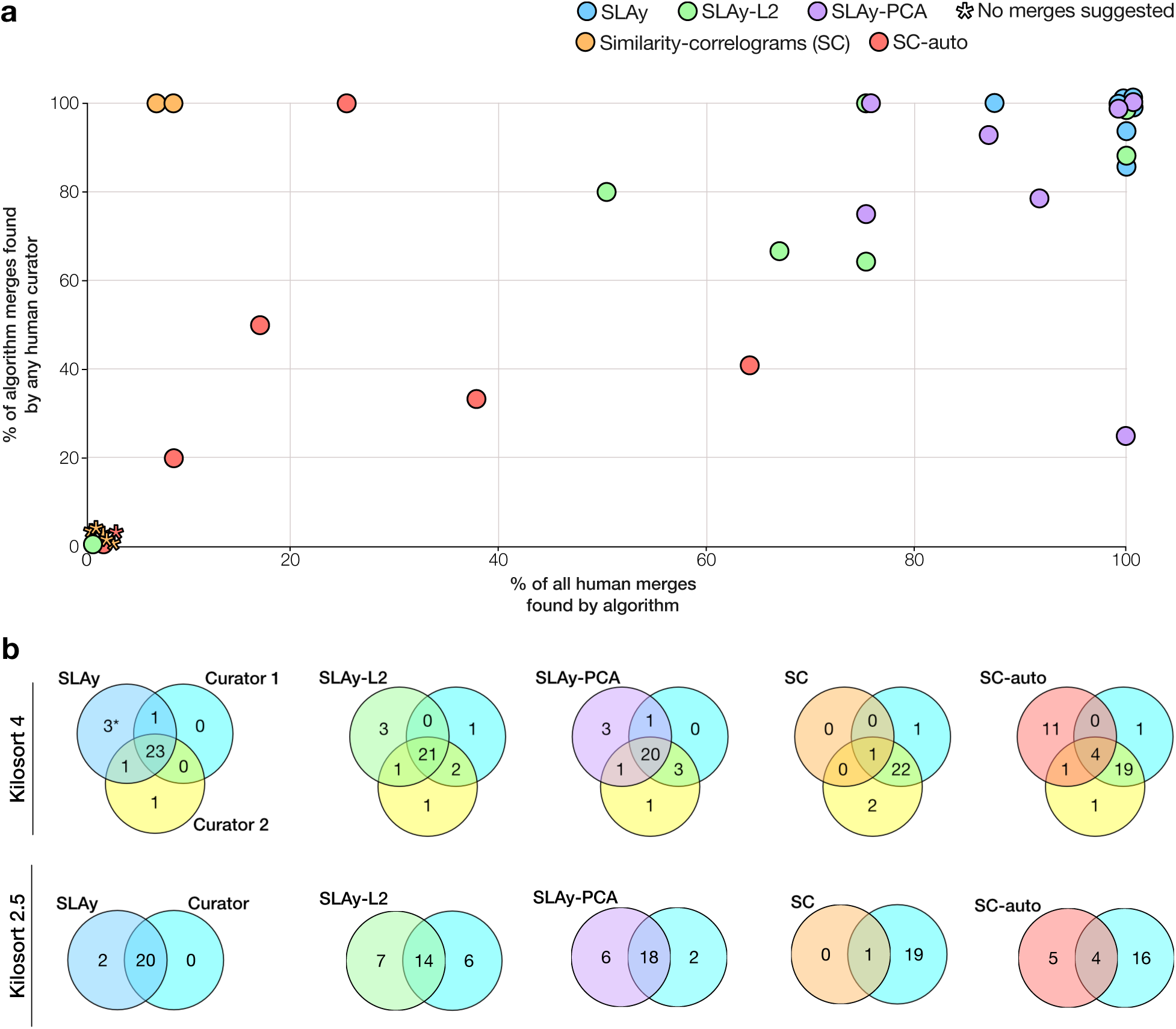
SLAy achieves high agreement with human curators on unaltered real datasets. **(a)** Precision and recall on curator-identified merges for each algorithm. Points at (0,0) and (100, 100) are jittered for visibility. KS4-Monkey midbrain is not shown, as neither the algorithms nor human curators identified any merges in that dataset. **(b)** Venn diagrams showing the overlap in merges between each algorithm and human curators, pooled across Kilosort versions.

Many of the merges identified by SLAy corresponded to common failure modes of template matching. For example, in the KS2.5-hippocampus recording, SLAy and both human curators merged together an oversplit bursting neuron despite a large difference in their mean waveforms (Fig. S8). The waveform similarity was not enough to result in a merge on its own, but the asymmetric peaked CCG, characteristic of a split bursting neuron, resulted in a high CCG structure metric that pushed the merge decision metric over the threshold. To illustrate the importance of merging for downstream analyses, we adapted a Hidden Markov model (HMM)-based algorithm [36] to detect bursts in spike trains (Fig. S14a). We applied our burst detection algorithm before and after performing merges identified by SLAy. For the insula and hippocampus recordings, SLAy’s identified merges resulted in a marked increase in number of detected bursts and burst intensity (Fig. S15).

Two merges in the KS4 mouse hippocampus dataset were identified by only SLAy and later approved by a human curator upon blinded re-evaluation (Fig. 6). Units within each merge corresponded to large temporal shifts of the same spike waveform (Fig. S9). The spikes detected with both or only one unit’s template did not differ visually, suggesting that these errors stemmed from improper template learning. The relative shifts in spike time identification resulted in a low L1 similarity between their mean waveforms; because human curators used a GUI that applies an L1 difference threshold to screen pairs of units, both curators missed these merges initially. Notably, these pairs did not pass the screening step even though curators used a more permissive threshold than usual (normalized L1 difference *<* 0.5 vs. default value of 0.25). Such curator misses thus may occur more frequently than observed here. Interestingly, both merges exhibited features of two types of template matching failures: a large peak at the center of the CCG, characteristic of units with co-detected spike times, and an asymmetric CCG with short, regular inter-spike intervals, characteristic of an oversplit bursting neuron. Further inspection of the spike times for the constituent units revealed that while many spikes were co-detected as part of both units, some spikes were assigned to only one unit or the other — with the assignment differing even for spikes within a single burst. This means that each oversplit unit contained unique information about the (presumed) ground-truth neuron, and thus merging them is vital to avoiding spurious conclusions about coding properties. That these merges were initially missed by both human curators highlights that, in addition to automating a tedious manual process, SLAy can improve the fidelity of neural datasets and downstream analyses.

A small number of merges were identified only by SLAy and not approved by a human curator upon blinded re-evaluation (Fig. S10). One such case involved a pair of high-amplitude units whose mean waveforms differed slightly but noticeably in the repolarization phase. Because this difference was relatively small compared to the overall magnitude of the spikes, it is likely that the autoencoder did not learn sufficiently distinct latent representations for spikes from these two units. Nonetheless, these rare false positives were outweighed by SLAy’s high precision and recall with respect to merges found by human curators.

The ablated versions of SLAy showed a lower degree of alignment with human curators than the autoencoder-based version and performed comparably to each other overall. Across datasets, SLAy-L2 found ∼ 78% of merges that were initially identified by at least one curator (Fig. 6b) while SLAy-PCA found ∼ 86%. Compared to SLAy, both ablated versions also made more merges that human curators disagreed with: human curators agreed with only ∼ 78% of merges identified by SLAy-L2 and ∼ 82% of merges identified by SLAy-PCA.

The discrepancy in human-curator alignment between SLAy-L2, SLAy-PCA and SLAy varied between datasets (Figs. 6a, S6, S7). For example, all variants recovered all merges identified by human curators on the KS4 monkey motor cortex dataset. However, on the KS4 mouse hippocampus dataset, SLAy-L2 missed 50% of the merges identified by at least 1 human curator, and missed both valid merges that were originally missed by human curators but found by the full SLAy algorithm. On the same dataset, SLAy-PCA found one of the two merges initially missed by human curators (Fig. S9a-c), but missed the other, (Fig. S9d-f) which had a larger timeshift. SLAy-PCA missed two merges found by at least one human curator, and also suggested an additional merge not found by either human curator or the original SLAy algorithm.

Both ablated versions were considerably worse than the autoencoder-based SLAy on some individual recordings. On the KS2.5 rat insula dataset, a human curator disagreed with 5 out of 14 merges identified by SLAy-L2, compared to only 2 out of 14 disagreements with merges identified by autoencoder-based SLAy. SLAy-PCA was much worse the SLAy and SLAy-L2 on the monkey midbrain recorded with KS2.5: SLAy-PCA found the single merge found by both human curators (and SLAy), but suggested three merges not found by SLAy or either human curator. SLAy-L2 missed the merge found by human curators, but only suggested one extraneous merge rather than three.

Closer examination of the disagreements between SLAy-L2 and human curators revealed two main types of errors. One type of error was the failure to merge units that had visually similar mean waveforms that were shifted temporally relative to each other (Fig. S11a-d). These errors likely occured because L2 similarity is highly sensitive to timeshifts, as it squares and sums sample-wise differences between mean waveforms. Notably, these merges were missed by SLAy-L2 despite having high enough similarity to pass the screening stage of the manual curation GUI, though human curators used a more permissive GUI threshold than normal. Naturally, SLAy-L2 also failed to identify the timeshifted oversplits (Fig. S9) that human curators initially missed. Another type of error was merging units with differing spatial decay profiles (Fig. S11e-f). Because L2 similarity depends only on pointwise differences, two units with differing spatial decay may have a moderately high L2 similarity so long as they have similar waveforms on their highest amplitude channels. In some cases, this may lead to erroneous merges. The presence of these errors highlights the advantages of the autoencoder-based waveform similarity metric; SLAy-L2 was unable to identify pairs of units with even small timeshifts, while SLAy found merges with shifts large enough that they were missed by human curators. Additionally, because SLAy’s autoencoder condenses the multichannel waveform into a low-dimensional latent vector and is trained to reconstruct input spikes, it must directly capture the spatial decay profile of each unit. The autoencoder thus better distinguishes units with similar peak channel waveforms but different spatial decay profiles.

Disagreements between SLAy-PCA and human curators were less stereotyped than for SLAy-L2, though some examples of the classes identified above still occurred. For example, SLAy-PCA also missed the timeshift merge in Fig. S11a-b. SLAy-PCA also missed a 3-way merge identified by both human curators, SLAy, and SLAy-L2 in the rat insula KS4 dataset (Fig S12a-c). SLAy-PCA merged together only two of the three units, with the unmerged unit having a slightly temporally shifted mean waveform relative to the other two. This indicates that PC features for individual spike waveforms are sensitive to timeshifts, perhaps more so than L2 similarity. SLAy-PCA also made some errors similar to the spatial decay false positive merges from SLAy-L2 (Fig. S12f). Other disagreements between SLAy-PCA and human curators seemed more idiosyncratic, such as SLAy-PCA not merging two units with very similar mean waveforms and low CCG collisions (Fig. S12d-e), or conversely, merging units with highly separable amplitude distributions (Fig. S12g). Together, the results from the ablated versions of SLAy showed lower alignment with human curators than the autoencoder version, suggesting that nonlinear waveform features are necessary to achieve effective automatic merging performance.

As on the held-out simulated oversplits, the similarity-correlograms algorithm was overly conservative on the original datasets. Across all eight datasets, SC made only two merges: one each in the KS4 and KS2.5 rat insula datasets. Although human curators agreed with both merges, they correspond to only ∼ 4% of the total number of merges made by human curators. This low rate of merging is expected considering that the SC algorithm is highly similar to post-sorting merge steps already implemented in both Kilosort versions. Consequently, this indicates that the primary existing automerging algorithm does not correct enough oversplits to replace manual curation.

SC-auto identified 20% of the merges made by human curators. Though this was an increase relative to SC, it remains considerably lower than both variants of SLAy. This modest increase came at the cost of considerable extraneous merging: human curators agreed with only 36% of the merges made by SC-auto across datasets. These extraneous merges likely stem from limitations in SC’s design, which requires waveform *and* ACG-CCG differences to be below certain thresholds. However, both these conditions are unlikely to be satisfied in non-random oversplitting, e.g., oversplitting of bursting neurons, which result in units with lower waveform similarity and asymmetric CCGs. As a result, successfully merging simulated burst and/or amplitude oversplits during the parameter tuning process may have resulted in thresholds that are too permissive. This idea is supported by the values of SC-auto parameters shown in Table 3, which are considerably higher than the SC default values (0.25 for waveforms and 0.16 for ACG-CCG). Importantly, this reflects a fundamental limitation of SC’s design: its reliance on simultaneous waveform and ACG-CCG similarity makes it ill-suited for identifying non-random oversplits with high sensitivity.

Together, these results demonstrate that the novel components of SLAy’s design —– specifically the use of autoencoder-based waveform similarity and the thresholding of a linear combination of metrics –— enable the automated correction of structured oversplitting with high human-curator alignment across a wide range of datasets. While automating and standardizing unit merging alleviates critical bottlenecks for throughput and reproducibility in large-scale ephys data processing, SLAy’s utility may extend further: by identifying valid merges that human curators miss, SLAy could improve the reliability of spike-sorted recordings and the resulting scientific findings.

## 3 Discussion

Automated spike sorting using template matching has become the dominant method for processing high-density ephys recordings. However, despite the inclusion of post-sorting merge steps, template matchers frequently introduce oversplitting errors that can compromise the validity of downstream analyses —– for example, by biasing estimates of correlation between pairs of neurons [23]. Correcting these errors is particularly critical when analyzing recordings susceptible to oversplitting, such as those from brain regions with bursting neurons or recordings characterized by significant probe motion. While these errors typically require labor-intensive manual correction, we introduced SLAy to identify and correct them in a quantitative and reproducible fashion. SLAy suggests valid merges across a variety of animal models, brain regions, probe geometries, and spike sorters. We demonstrated that performing these merges improves the validity of downstream analyses; specifically, SLAy merged oversplit bursting neurons in CA1 and the insula, accurately assigning spikes within a burst to a single neuron and resulting in an increase in both the number and intensity of detected bursts.

Using SLAy across the wide range of datasets shown here may require parameter tuning. Many parameters, specifically those involved in the calculation of the individual SLAy metrics (e.g. CCG bin size, refractory period width), can be set to standardized values for each animal model or brain region. For the remaining parameters, namely the coefficient for the CCG structure metric (*k*_1_) and the merging threshold on the final metric, we developed an automatic parameter tuning procedure that adapts to the specific characteristics of each dataset. Our parameter setting scheme introduces structured simulated oversplitting that mimics the failure modes of template matching algorithms. We were initially concerned that this might result in the selection of overly conservative parameters, as the spikes in our simulated oversplits were already grouped into a single unit by the spike sorter, and thus may be “easier” to detect than real oversplits. However, we show in practice that SLAy detected all oversplits identified by multiple curators, indicating that the simulated oversplits do yield SLAy metrics similar to those of real oversplits. The fact that Kilosort oversplits some presumed ground-truth neurons but not others (i.e. bursting neurons that we artificially oversplit), may be related to the inherent stochasticity in the greedy spike detection process. Nevertheless, we show our automated procedure result in high alignment with human curators, removing the need for manual adjustment of parameters. Our codebase, available as a pip-installable package, retains support for manual parameter adjustment for users who want to align SLAy’s merges more closely with their own needs. Depending on the considerations for downstream analyses, users may also want to manually review merges suggested by SLAy rather than accepting them automatically. For example, SLAy corrects a template matching failure mode where the simultaneous firing of different neuronal subparts, which are recorded on noncontiguous parts of the probe, are split into separate units. Although the codetection merges suggested by SLAy have high waveform similarity and very large peaks at the centers of the CCG, suggesting that merged subparts almost always fire together, some scientific questions may require analysis of the firing of different neuronal compartments. The full version of SLAy, including the autoencoder-based waveform similarity and automatic parameter selection, can thus run in a semi-automated mode that outputs a list of suggestions for manual review. This approach still saves time compared to traditional manual curation, since users would have to review a much smaller list of candidates compared to all pairs of clusters with moderately similar waveforms.

Although few postprocessing tools exist to automatically correct spike sorting output, with none widely adopted, similar procedures have been devised for other goals. For example, [37] and [38] use mean waveform metrics and spatial localization of neurons among other metrics to track individual neurons across recordings spanning multiple days. These approaches, though not designed for post-sorting correction of a single spike sorter’s output, share conceptual similarities with SLAy’s methodology.

To our knowledge, SLAy represents the first use of a neural network to define a feature space specifically for calculating spike waveform similarity. While other studies have utilized autoencoders in electrophysiology, their applications differ significantly. For instance, Beau et al. use the latent space of an autoencoder to predict neuronal cell type, but their model only encodes the peak-channel mean waveform [39]. Information regarding the spatial decay of recorded spikes, while perhaps secondary for cell-type classification, is essential for determining whether two putative units correspond to the same ground-truth neuron. Other approaches, such as Yet Another Spike Sorter (YASS) [19], use autoencoders to denoise waveforms before clustering; however, since YASS is trained on synthetic data without realistic noise, it can hallucinate spikes in noise-only snippets and does not utilize the latent vectors as a comparative feature space. Similarly, DeepInterpolation [40] employs a convolutional autoencoder for denoising but uses residual connections that allow information to bypass the bottleneck, making the resulting encoding unsuitable for feature extraction.

Given that many modern spike sorters still rely on per-channel principal components (PCs) as features [17, 18], the advantages demonstrated by our autoencoder-based features — namely higher explained variance, robustness to jitter in spike-time identification, and more accurate representation of spatial decay — suggest a promising new direction. Future research should investigate whether clustering directly within an autoencoder-based feature space can improve the accuracy of spike-sorting algorithms, potentially reducing the burden of post-hoc correction.

Although it is clear that oversplitting can bias single-neuron analyses, it may be argued that population analysis techniques involving dimensionality reduction are robust to oversplitting. In this case, one might assume that split units would be sufficiently correlated to collapse together when performing dimensionality reduction. However, this assumption holds only if oversplitting is completely unbiased, which is generally not true. While SLAy rarely suggests merges that human curators disagree with, previous studies have shown that overmerging biases downstream analysis only if the merged units have significantly anticorrelated firing [41].

SLAy represents a significant step towards fully automating spike sorting curation – a critical unmet need given the growing scale of electrophysiological recordings. SLAy-L2 has already been integrated into SpikeInterface, and the full version of SLAy, featuring autoencoder-based waveform similarity and automatic parameter tuning, is available as a pip-installable package. The standalone package also supports SpikeInterface objects, making SLAy applicable across a wide range of recording hardware and spike sorters, and compatible with existing data processing pipelines. SLAy ran in *<* 10 minutes for the largest recording we tested (410 single-units, 60-minute recording, benchmarked on an NVIDIA RTX 3080), more than an order of magnitude faster than manual curation.

Code: https://github.com/saikoukunt/SLAy

## 4 Methods

### 4.1 Recordings

We applied SLAy to 4 Neuropixels recordings:

- Mouse hippocampus, Neuropixels 2.0: This recording was collected by Dohoung Kim in Albert Lee’s lab at the Janelia Research Campus. It was about 60 minutes long across 384 channels.
- Male Long-Evans insula, Neuropixels 1.0 during a free drinking task: This recording was 60 minutes long across 384 channels, and was collected by Aamna Lawrence in Patricia Janak’s lab at Johns Hopkins University.
- Female macaque midbrain, 128-channel prototype Neuropixels NHP during vestibular stimulation (using an Intan recording system): This recording was 10 minutes long across 128 channels, and was collected by Robyn Mildren in Kathleen Cullen’s lab at Johns Hopkins University.
- Macaque motor and premotor cortex, Neuropixels 1.0: This recording is a publicly available recording from Bijan Pesaran’s lab [31]. It was 10 minutes long across 384 channels.

### 4.2 Processing and spike sorting

For the mouse hippocampus, rat insula, and macaque motor cortex recordings, preprocessing and spike sorting were performed using CatGT and SpikeInterface. Common average referencing, bandpass filtering, and artifact removal were performed using CatGT. Preprocessing for the macaque midbrain data was done using a custom Python script, due to incompatibility of CatGT and Intan recording hardware. We then used SpikeInterface to spike sort each recording with Kilosort 2.5 and Kilosort 4 and identify single-units using quality metric filters.

### 4.3 Unit quality labeling

After sorting, we used a custom automated pipeline to automatically label units as single-unit, multi-unit, or noise. units were marked as noise if they failed any of the following quality metric tests:

- firing rate *>* 0.1 Hz
- signal-to-noise ratio *>* 2,

The remaining units were marked as multi-unit if they had sliding refractory violations *<* 0.1, and single-unit otherwise.

### 4.4 Autoencoder architecture and training

#### Architecture

The encoder contains 7 layers – a flattening layer that converts the 8 channels by 40 samples snippet into a 320 dimensional vector, followed by 3 pairs of dense linear + Gaussian error linear unit (GeLu) activation layers. The layer sizes are [320 → 600, 600 → 300, 300 → 15], which produces a 15-dimensional feature vector. The decoder also contained 7 layers, in reverse order – 3 pairs of dense linear+ Gaussian error linear unit (GeLu) activation with hidden unit counts [15 → 300, 300 → 600, 600 → 320] followed by an unflattening layer that reshapes the snippet to the original 8 channels by 40 samples shape.

#### Training

We trained a separate network for each dataset. Spike snippets for autoencoder training were extracted from the preprocessed recordings, and consisted of the peak + 7 nearest channels, using a window from 10 samples before to 30 samples after the inferred spike time. We extracted a maximum of 1000 snippets per unit, and units with less than 100 total spikes were ignored. Snippets were split into training and testing sets using an 80-20 split, and only the training set was used for model training. The autoencoder training procedure uses an L2 loss function, a batch size of 128, a learning rate of 0.001, and 25 epochs of training for all networks. All networks were implemented and trained using PyTorch and CUDA on an NVIDIA 3080 RTX GPU.

### 4.5 Waveform similarity calculation

To calculate the waveform similarities between units, we first calculate a spike feature vector for each spike snippet and take the mean feature vector of each Kilosort unit as the unit centroids. For each pair of units, we then calculate the unnormalized Euclidean distance between unit centroids: **dist_ae_** = ||*µ*_1_ − *µ*_2_|| where *µ*_1_ and *µ*_2_ are the unit centroids. We normalize these distances by the standard deviations of the units, so that the similarity metric is invariant to distortions in latent space geometry. The waveform similarity is thus calculated as

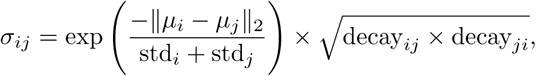

Exponentiating the negative distance yields a similarity metric between 0 and 1 rather than a distance metric. This metric measures the shape similarity between two waveforms but does not include spatial information. To re-introduce spatial information and penalize similarity between waveforms that are similar but on different parts of the probe, we calculate a spatial decay between peak channels for each unit:

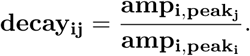

If the peak channels are far apart, *amp_i,peak__j_* and *decay_ij_* will be close to 0. If the peak channels are the same (*amp_i,peak__j_* = *amp_i,peak__i_*), or the two units have different, nearby peak channels due to noise but decay very little across channels, the decay metric will be close to 1. To calculate the final metric, we penalize the distance-based similarity using the calculated decay metrics:

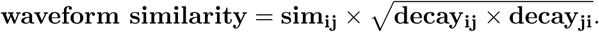

After calculating the waveform similarities, only unit pairs with a similarity greater than a threshold (0.4 by default) are considered for merging, and the rest are assumed to not be oversplit. This filtering step considerably improves the runtime of SLAy by avoiding computationally expensive CCG calculations for dissimilar units.

### 4.6 Cross-correlogram significance metric

Cross-correlograms were computed using a multithreaded Python function with a window size of 50 milliseconds and a minimum bin width of 1 ms. To determine CCG significance, we first selected a window of interest by applying a trough finding algorithm to a low-pass filtered version of the second derivative of the CCG. The identified troughs in the second derivative correspond to the locations of sharp changes in the CCG, so the window of interest was taken as 1.5 times the distance from the CCG center to the most central set of troughs. If the CCG did not have a high enough event rate in the window of interest (defined by a user-set threshold), the window was expanded to the 1.5 times the subsequent trough location until the CCG had an acceptable event rate in the window of interest. We then calculated the CCG significance metric as the earth mover’s distance between the windowed CCG and a uniform distribution divided by a scaling factor of 0.25. 0.25 corresponds to the unnormalized EMD between a single peak at the center of the CCG and a uniform distribution, which provides a reasonable upper bound on the EMD for a unit pair. We thus divided by 0.25 to normalize the metric to be between 0 and 1. If the window based on the furthest set of troughs still had an event rate that was too low, we penalized the metric by the squared ratio between the observed event rate and the minimum allowed event rate. Window sizes and bin widths specified above are the default values, but all values are user-adjustable.

### 4.7 Refractory period violation calculation

Refractory period violations are computed using 2-second window CCGs calculated in the same manner as above. Candidate refractory periods ranged from 1 ms to 10 ms (user-adjustable). We first calculated the maximum allowed refractory period contamination. We estimated the baseline CCG event rate as the maximum of the low-pass filtered CCG, and took the maximum allowed event rate to be a fraction (0.25 by default) of the baseline. For each candidate refractory period, we calculated the observed number of refractory period violations and the maximum allowed number of refractory period violations. We then took the refractory period penalty to be the cumulative probability of observing up to the observed number of violations under a Poisson distribution with a mean equal to the maximum allowed violations. If the observed number of violations is small compared to the allowed violations, the penalty will be small. If the observed number of violations is large compared to the allowed violations, the penalty will be large.

#### Algorithm 1

SLAy merging algorithm

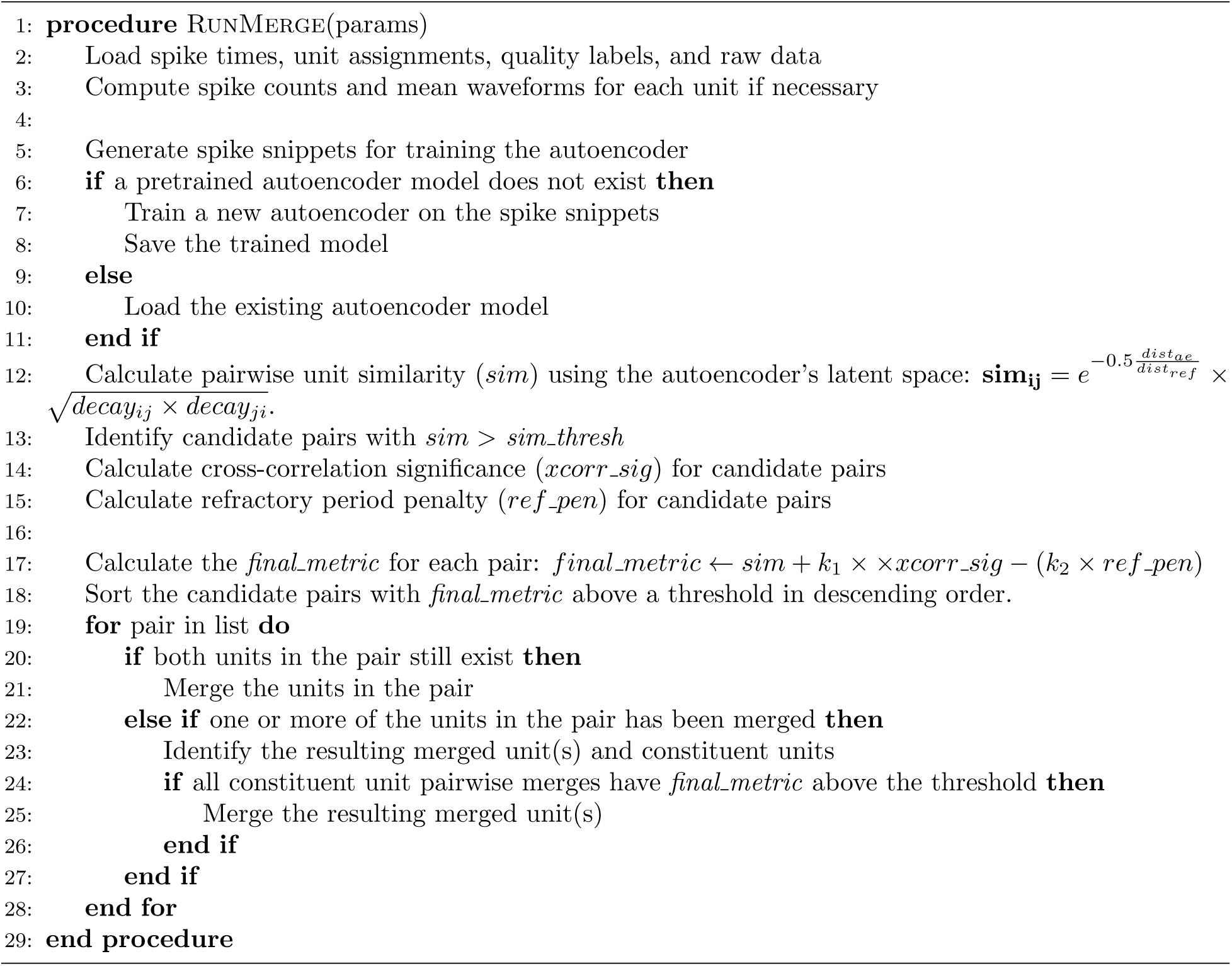

### 4.8 Merging process

Our final merge decision metric, M*_ij_*, integrates the above metrics to determine whether to merge units:

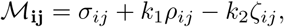

where *σ_ij_, ρ_ij_*, and *ζ_ij_* are the waveform similarity metric, cross-correlation significance metric, and refractory period penalty, respectively. *k*_1_ and *k*_2_ are coefficients ≤ 1 that control the relative importance of the metrics. To perform merges, we sort the unit pairs by M*_ij_* in descending order and iterate over them. If M*_ij_* is greater than a threshold and both units in the pair still exist, we merge them. If M*_ij_*is greater than the threshold but one of the units has already been merged into a new unit, we have a potential multi-way merge. We perform the multi-way merge only if all pairwise merges between constituent units have an M*_ij_* greater than the threshold.

### 4.9 Burst detection

To detect bursts in spike trains, we developed the EM-Kleinberg model, an extension of a model used to identify bursts in text streams [36]. The original model is a generative hidden Markov model that assumes the system is in one of a few hidden states, each of which is associated with Poisson emissions at a certain firing rate. Each HMM state *i_k_* is parameterized by a Poisson rate *a_k_* ≥ *a*_0_*s^k^*that governs the distribution of ISIs, where *a*_0_ is the neuron’s overall average firing rate and *s >* 1 is a multiplicative scaling factor that controls how much the firing rate increases between HMM states. The “states” here are simply a modeling choice to adequately capture the statistics of the observed ISIs, and do not necessarily correspond to actual biophysical states of a neuron. We use a method from the original algorithm [36] to pre-calculate the number of HMM states based on *s*, the length of the spike train, and minimum observed ISI.

A period of intensity *k* is defined as an interval where the neuron’s state is ≥ *i_k_*. Transitioning to less active states (i.e. lower intraburst firing rate) has no cost, while transitioning to a more active state has a cost proportional to the number of skipped states. The state sequence is inferred by minimizing a cost function that includes the transition cost and the negative log-likelihood of the observed ISI given the underlying state. The bursts are then identified by designating states above a user-defined firing rate as bursting states. We find that initializing with *s* = 3 and designating firing rates higher than 50*Hz* works well for neuronal spike trains. The original algorithm fixes the Poisson rates at *a_k_* = *a*_0_*s^k^*, while we initialize the rates using *s* but then estimate the rates during optimization using expectation-maximization (EM), with the constraint *a_k_* ≥ *a*_0_*s^k^*.

The first step of the EM process is similar to the optimization in the original model; it uses the forward-backward algorithm to estimate the optimal state sequence given the firing rates of each state and the observed ISIs. The M step then uses a maximum likelihood estimator to update the estimated firing rates of each state given the inferred state sequence. Although it is typical to run EM-algorithms until the likelihood converges, we instead minimize runtime by stopping inference after 5 iterations.

To assess the performance of our burst detection algorithm, we generated simulated spike trains with a wide variety of firing statistics [42] (Fig. S14c) and compared our algorithm’s performance against those of three existing methods: MaxInterval [43], logISI [44], and the original HMM algorithm [36]. The Kleinberg and EM-Kleinberg algorithms were implemented in Python using parameters *s* = 5*, γ* = .3, designating states ≥ 2 as bursty states. On spike trains consisting of only bursts, our algorithm achieves near-perfect classification accuracy (burst vs. non-burst) regardless of the burst length or intra-burst frequency (Fig. S14b). The MaxInterval and logISI methods achieve near-perfect classification on two out of three of the simulated spike trains, while the the original Kleinberg HMM algorithm achieves near-perfect accuracy on one simulation and moderate accuracy on the other two. Our algorithm also outperforms the others on simulated spike trains containing both bursts and single spikes.

We applied this burst detection algorithm before and after automatic merges performed by SLAy, to illustrate the importance of unit merging for downstream analysis.

### 4.10 Simulated oversplitting

For each of the four error types described below, we first identified a pool of suitable candidate units and then randomly selected a number of these candidates (*num splits*) to be split. The ratio of each type of split was equal. To ensure that the split units matched the quality of existing single-units, we re-applied single-unit quality filters to the candidate splits, and only performed the splits that would result in units that passed all quality filters. Because of this, the percent of units split differs between datasets. To generate error bars for recall and percent of total units merged, this entire procedure was repeated 5 times for each dataset.

#### Burst splitting

To simulate oversplitting due to bursting, we first identified candidate units where at least 30% of their inter-spike intervals (ISIs) were less than 20 ms. Within these units, we defined a burst as a sequence of 3 or more spikes with ISIs under 20 ms. To ensure the resulting units were sufficiently large for analysis, only units where the total number of bursting spikes was greater than 600 were included as candidates. For each selected unit, we split it into two new units: the first contained all non-bursting spikes and the spikes from the first half of each burst, while the second contained only the spikes from the second half of each burst.

#### Amplitude splitting

Earlier versions of Kilosort penalized the variance of the amplitude distribution of each unit and thus oversplit units with highly variable spike amplitudes. To model errors from high amplitude variance, we identified candidate units whose amplitude variance was between the 75th and 95th percentile of the distribution for that recording. This allowed us to target high-variance units while excluding extreme outliers that may have resulted from over-merging errors. For each selected unit, we chose a random *split ratio* between 30% and 50% and assigned the *split ratio* of spikes with the highest amplitudes to a new unit.

#### Drift splitting

To simulate oversplitting due to probe drift, we constructed a time-dependent splitting probability defined by a three-part piecewise linear function. The probability of being assigned to the new unit ramped from 0 to 0.1 over the first 40% of the recording, from 0.1 to 0.9 over the middle 20%, and from 0.9 to 1.0 over the final 40%. As candidates for this split, we selected units that were active throughout the recording, defined as having at least 30% of their spikes in each half of the recording. For each selected unit, we iterated through its spikes and assigned a spike to the new unit if a randomly sampled number was less than the splitting probability at that spike’s time.

#### Random splitting

To simulate random oversplitting, we marked all units that passed the noise and multi-unit criteria as candidates. For each unit selected to be oversplit, we selected a random *split ratio* of spikes between 30% and 50% and randomly assigned that percentage of spikes to a new unit.

### 4.11 Automatic parameter selection

The SLAy algorithm incorporates the automatic parameter selection scheme described in Section 2.3.2. We fixed *k*2 = 1 and evaluated the recall and percent of total units merged for the parameter grid (i.e. all possible combinations of each parameter) over *k*1 and the threshold for the final metric:

- *k*1 : 0.25 to 0.55, in steps of 0.05.
- merge threshold: 0 to 0.85, in steps of 0.05

We also applied the automatic parameter selection to the similarity-correlograms algorithm, searching over a grid of thresholds for waveform difference and ACG-CCG difference:

- waveform difference threshold: 0.0 to 1.0, in steps of 0.05
- ACG-CCG difference threshold: 0.0 to 1.0, in steps of 0.05

The default parameters in SpikeInterface for the similarity-correlograms are waveform difference threshold = 0.25 and ACG-CCG difference threshold = 0.16.

Tables 1-3 show the parameter selected for each algorithm on each dataset.

**Table 1:**
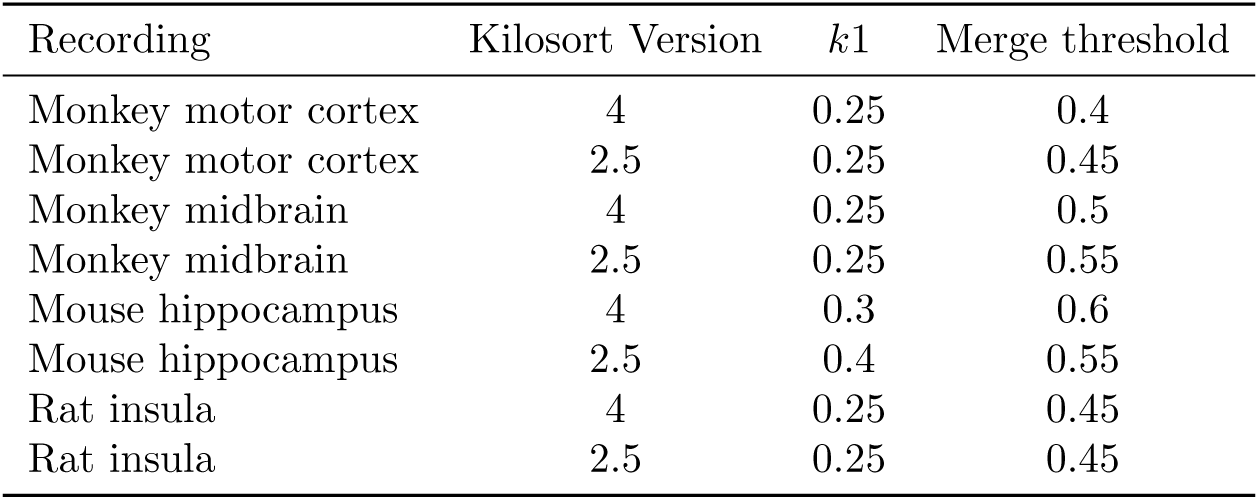
Autoselected parameters for SLAy on each dataset.

**Table 2:**
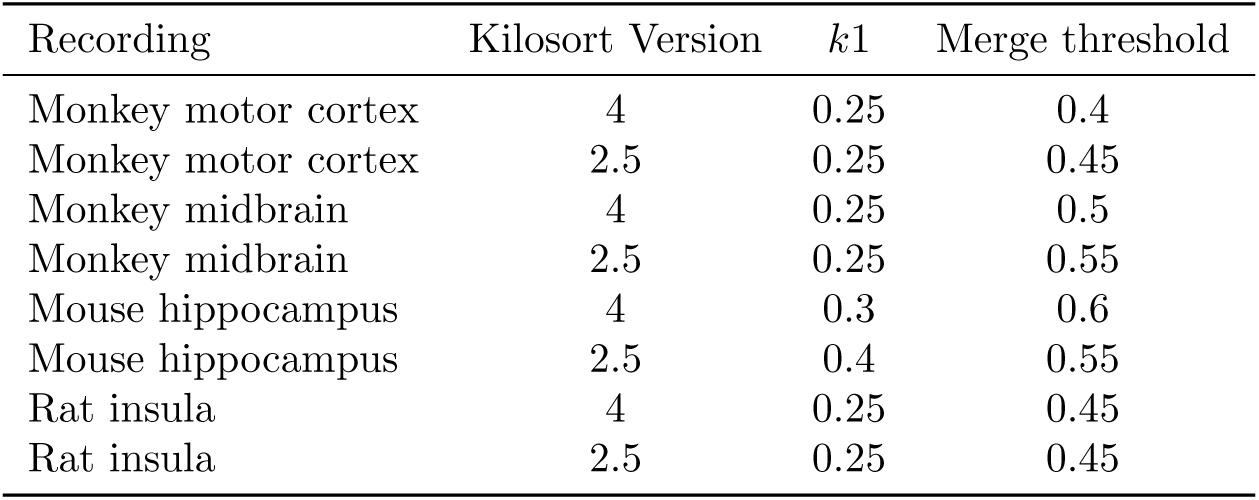
Autoselected parameters for SLAy-L2 on each dataset.

**Table 3:**
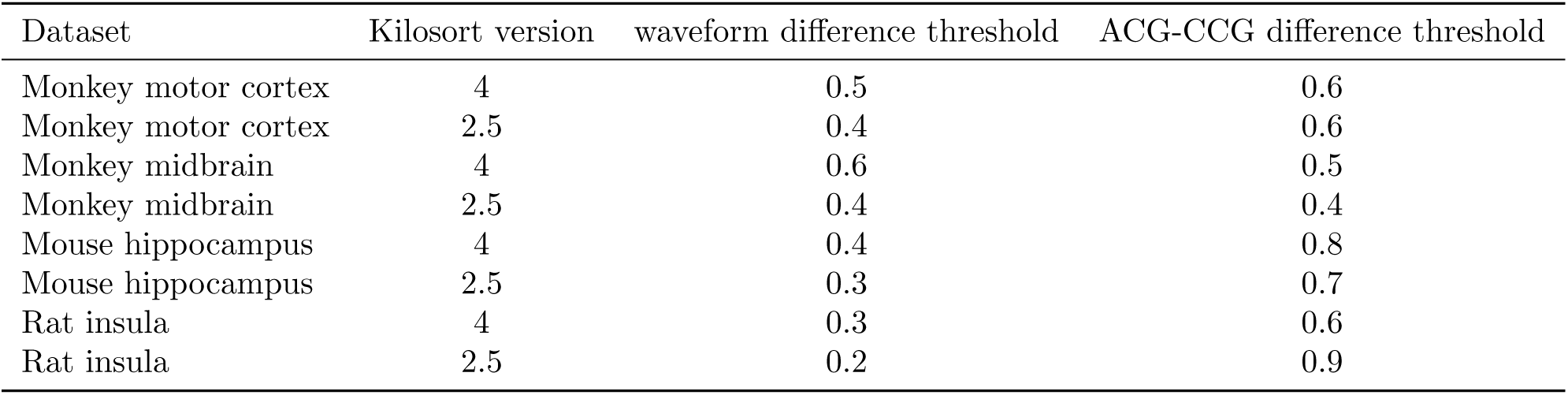
Autoselected parameters for similarity-correlograms on each dataset.

### 4.12 Testing sensitivity to autoencoder and simulated oversplitting random seeds

To assess SLAy’s sensitivity to randomness in both autoencoder training and the simulated oversplits used for automatic parameter selection, we ran SLAy on two datasets, the monkey motor cortex sorted with KS2.5 and with KS4. For each dataset, we trained autoencoder’s using 5 different random seeds—this affected both the initial weights of the autoencoder and the dataloader used during training, i.e. the sequence of training examples during training. For each pair of different autoencoders on each dataset, we calculated the Pearson’s correlation between the pairwise unit similarities computed using each autoencoder’s latent space. These are reported in Tables S1 and S2.

We then tested the sensitivity of the autoselected parameters to the random seed used in artificial split generation, which affects which units are split, the type of split applied to each unit, and for the random and drift splits, which new unit each spike in a unit was assigned to. For each of the 5 autoencoders above, we ran SLAy with 5 different simulated oversplitting seeds, for a total of 25 different runs per dataset. The automatically selected parameters for each run are reported in Tables S3 and S4. We also examined the variability of the suggested merges across runs. For both datasets tested, all 25 runs produced the exact same set of merges.

### 4.13 Significance testing for performance testing on simulated oversplitting

For the statistical significance of the performance on simulated oversplitting reported in Section 2.4, we averaged the recall across draws for each dataset to get a vector of 8 mean recalls for each method (one mean recall per recording-sorter combination). We performed one-sided paired t-tests for each pair of methods (SLAy *>* SC-auto, SLAy *>* SC, SC-auto *>* SC) with Holm-Bonferroni correction for multiple comparisons. We verified that the distribution of differences for each pair of methods was symmetric without outliers, making the paired t-test suitable. Effect sizes reported were calculated using Cohen’s *d*.

### 4.14 Accuracy and Validation Metrics

Manual curation was performed in the SpikeInterface GUI, based on the guidelines in the Phy practical guide to manual curation [33]. To avoid considering every single pair of units, the SpikeInterface GUI’s merge panel was used to pre-select units with less than 0.5 normalized L1 difference between mean waveforms. Only pairs under this threshold were considered further for merging. This general procedure of inspecting only pairs with waveform similarity above a certain threshold is standard practice in manual curation.

To quantify the level of spatial template mismatch for a unit, we calculated the distance between the centroids of the mean and template waveforms. Centroids were calculated as an amplitude-weighted average of the positions of channels where the waveform amplitude was at least 20% the peak amplitude.

To quantify the change in burst intensity and number of bursts for a suggested merge, we run the EM-Kleinberg burst detection algorithm on the constituent unit spike trains before merging, and the merged spike train after merging. The change in burst intensity was calculated as the percentage change in average HMM state during bursts.

#### Code Availability

Code and documentation for SLAy is available at

## Acknowledgements

We thank Dohoung Kim, Albert Lee, and Patricia Janak for allowing us to use their Neuropixels recordings to test and validate SLAy, as well as helpful discussions and feedback.

SK, AC, AG, and TH were supported by NIH grant NIH U01NS115587. ASC was partially supported by NSF CAREER award 2340338. AL was supported by NIH R01AA031609 and NIH R01AA027213. RM was supported by the Kavli Neuroscience Discovery Institute Distinguished Postdoctoral Fellowship and the Natural Sciences and Engineering Postdoctoral Fellowship. This work was supported by the National Institutes of Health/National Institute on Deafness and Other Communication Disorders (R01-DC002390 and R01-DC018061) to K.E.C.

## Supplementary Material

**Figure S1:**
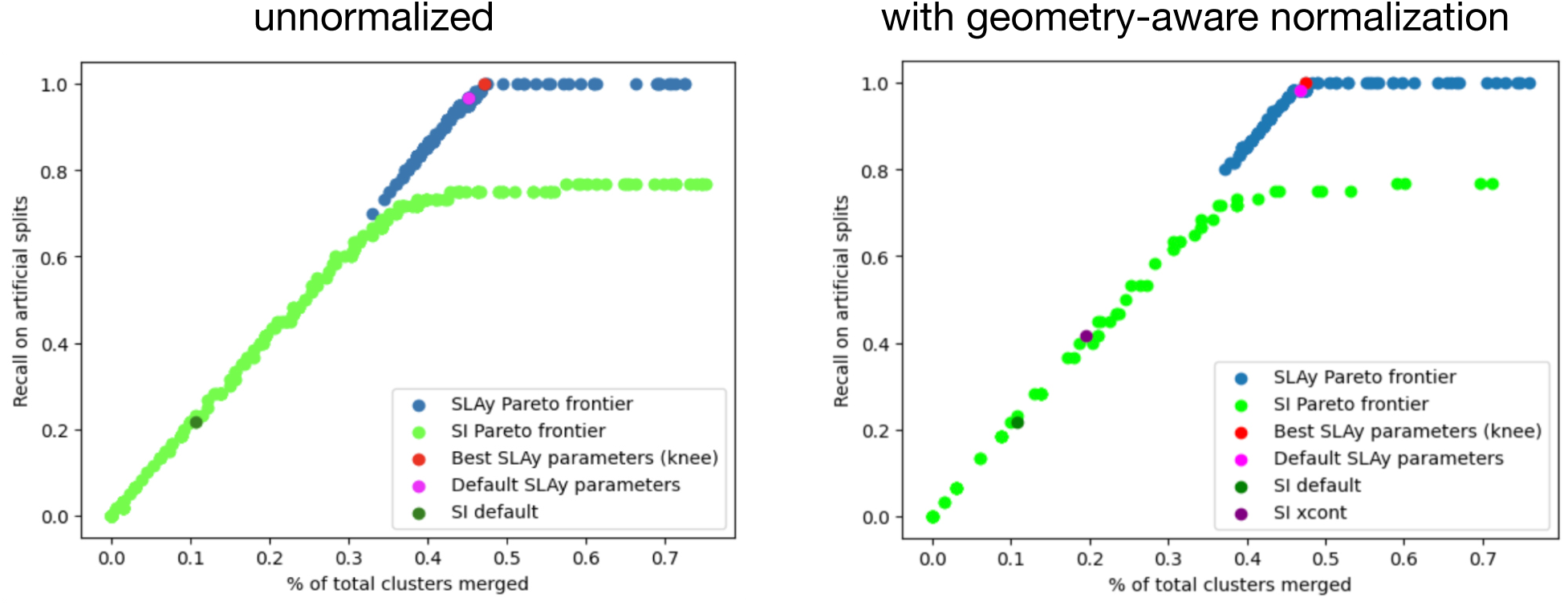
Geometry-aware normalization of waveform similarity metric reduces SLAy’s sensitivity to parameters. **(left)** Parameter sweep for SLAy without no normalization (blue). **(right)** Parameter sweep for SLAy with geometry-aware normalization for the same parameter grid as on (left). Compared to the unnormalized case, the left side of the curve starts about 0.1 higher, indicating that the geometry-aware normalization allows “bad” parameter combinations to achieve improved performance. Additionally the autoselected (pink) and default (red) parameters are closer in performance than the unnormalized case. This suggests that the normalization reduces disparities in performance between different parameter sets — i.e. it makes SLAy less sensitive to parameter selection.

**Table S1:**
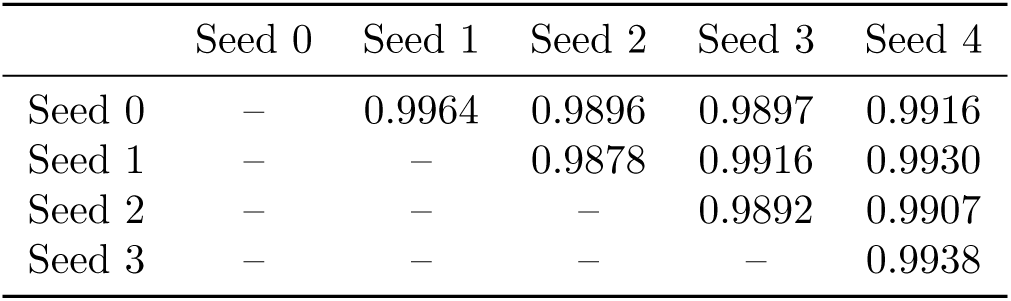
Unit similarity correlations (Pearson’s *r*) between autoencoder training seeds for monkey motor cortex KS4 dataset.

**Table S2:** Unit similarity correlations (Pearson’s *r*) between autoencoder training seeds for monkey motor cortex KS2.5 dataset.

|  | Seed 0 | Seed 1 | Seed 2 | Seed 3 | Seed 4 |
| --- | --- | --- | --- | --- | --- |
| Seed 0 | — | 0.9792 | 0.9845 | 0.9679 | 0.9694 |
| Seed 1 | — | — | 0.9916 | 0.9914 | 0.9887 |
| Seed 2 | — | — | — | 0.9863 | 0.9813 |
| Seed 3 | — | — | — | — | 0.9898 |

**Figure S2:**
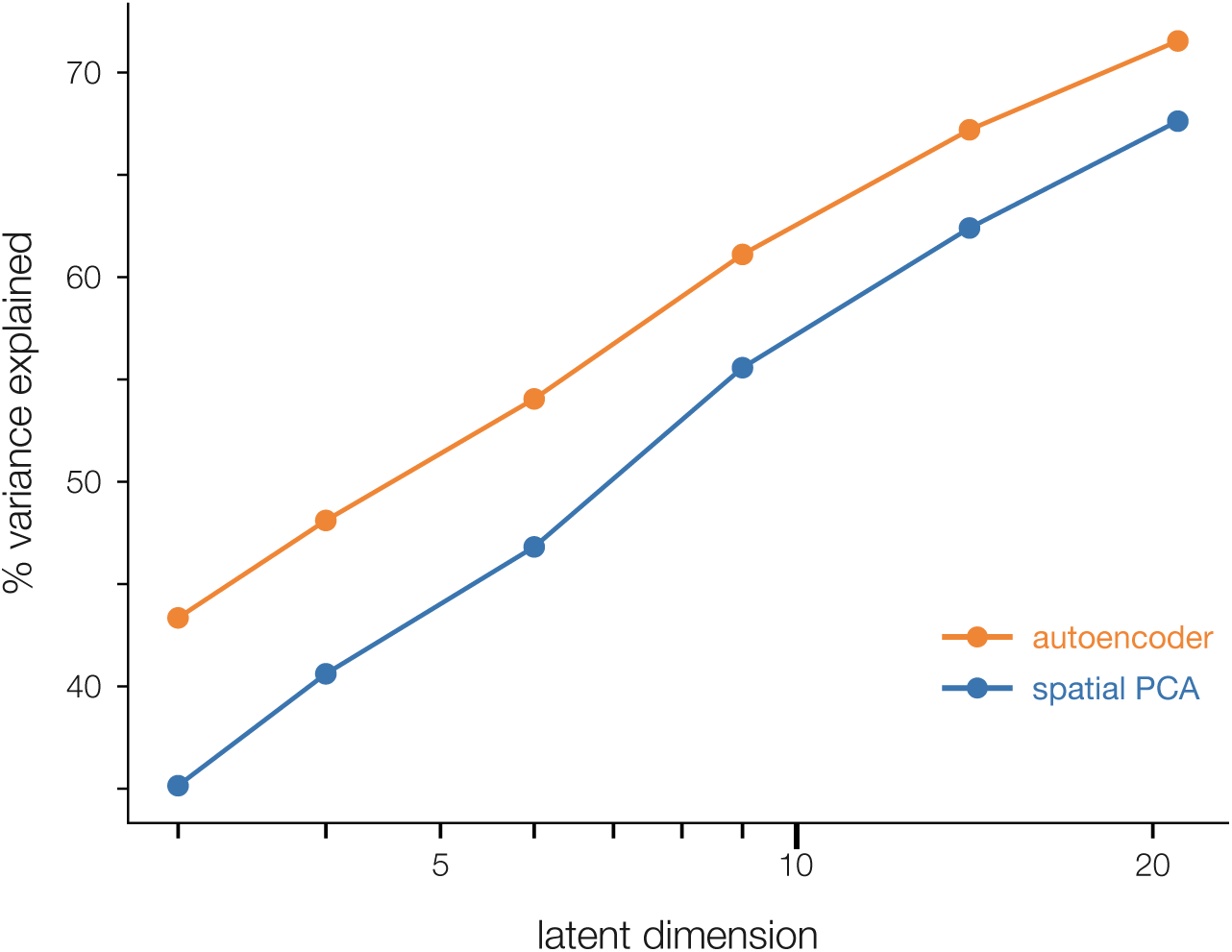
Autoencoder-based features explain more variance in waveforms than PCA-based features. For the same dimensionality, autoencoder-based features explain 5-10 % more variance in waveform shape than PCA-based features.

**Table S3:** Automatically selected parameters on KS4 monkey motor cortex dataset for combinations of autoencoder and simulated oversplitting seeds.

|  | AS Seed 0 |  | AS Seed 1 |  | AS Seed 2 |  | AS Seed 3 |  | AS Seed 4 |  |
| --- | --- | --- | --- | --- | --- | --- | --- | --- | --- | --- |
| | $k_1$ | thresh. | $k_1$ | thresh. | $k_1$ | thresh. | $k_1$ | thresh. | $k_1$ | thresh. |
| AE Seed 0 | 0.25 | 0.4 | 0.25 | 0.4 | 0.25 | 0.4 | 0.25 | 0.4 | 0.25 | 0.4 |
| AE Seed 1 | 0.3 | 0.4 | 0.25 | 0.4 | 0.3 | 0.4 | 0.25 | 0.4 | 0.25 | 0.4 |
| AE Seed 2 | 0.25 | 0.4 | 0.25 | 0.4 | 0.25 | 0.4 | 0.25 | 0.4 | 0.25 | 0.4 |
| AE Seed 3 | 0.35 | 0.4 | 0.25 | 0.4 | 0.25 | 0.4 | 0.35 | 0.4 | 0.25 | 0.4 |
| AE Seed 4 | 0.25 | 0.4 | 0.25 | 0.4 | 0.25 | 0.4 | 0.25 | 0.4 | 0.25 | 0.4 |

**Table S4:** Automatically selected parameters on KS2.5 monkey motor cortex dataset for combinations of autoencoder and simulated oversplitting seeds.

|  | AS Seed 0 |  | AS Seed 1 |  | AS Seed 2 |  | AS Seed 3 |  | AS Seed 4 |  |
| --- | --- | --- | --- | --- | --- | --- | --- | --- | --- | --- |
| | $k_1$ | thresh. | $k_1$ | thresh. | $k_1$ | thresh. | $k_1$ | thresh. | $k_1$ | thresh. |
| AE Seed 0 | 0.25 | 0.4 | 0.25 | 0.4 | 0.25 | 0.4 | 0.25 | 0.4 | 0.25 | 0.4 |
| AE Seed 1 | 0.25 | 0.4 | 0.25 | 0.4 | 0.25 | 0.4 | 0.25 | 0.4 | 0.25 | 0.4 |
| AE Seed 2 | 0.25 | 0.4 | 0.25 | 0.4 | 0.25 | 0.4 | 0.25 | 0.4 | 0.25 | 0.4 |
| AE Seed 3 | 0.25 | 0.4 | 0.25 | 0.4 | 0.25 | 0.4 | 0.25 | 0.4 | 0.25 | 0.4 |
| AE Seed 4 | 0.25 | 0.4 | 0.25 | 0.4 | 0.25 | 0.4 | 0.25 | 0.4 | 0.25 | 0.4 |

**Figure S3:**
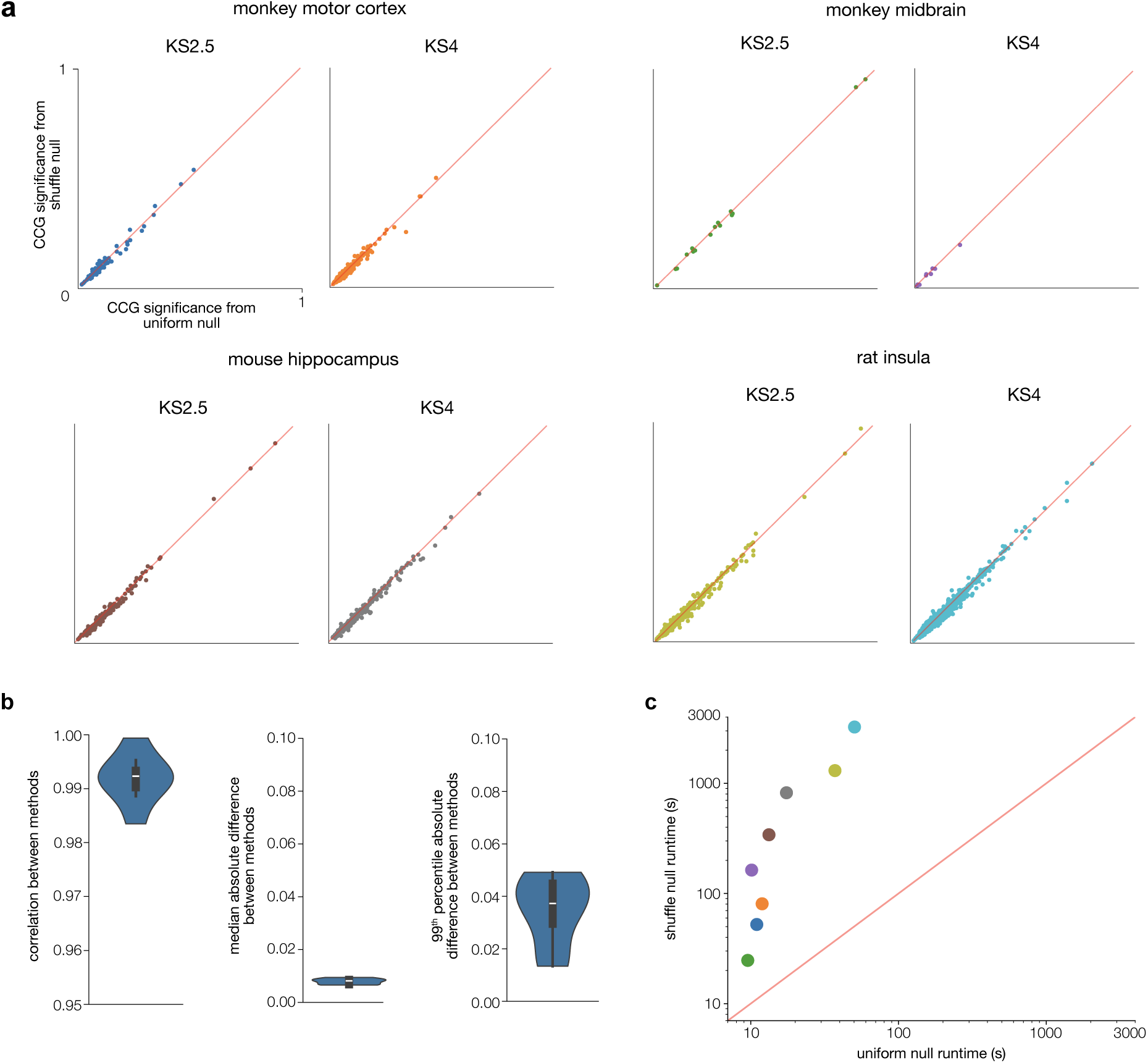
Uniform null vs. per-unit shuffle null for CCG significance metric. Using a uniform distribution as the null produces highly similar metric values while being much faster than a shuffle distribution. **(a)** Both methods of calculating null CCGs produce similar metrics across all recordings. Each colored dot represents a candidate unit pair for merging, and the red line represents *y* = *x*. **(b)** Mean correlation across recordings between null methods is high (0.992) and mean absolute differences are low (∼ 0.01 for median abs. difference, ∼ 0.04 for 99th percentile.) Note that the CCG significance metric is further multipled by a constant *<* 1, (0.25 by default) when calculating the final merging metric, so differences at this scale will have little impact on merging decisions. **(c)** The uniform null is about an order of magnitude faster to calculate than the shuffle null. For long recordings (∼ 1 hour) or recordings with many units (∼ 1000), the uniform null takes ∼ 100 s while the shuflle null takes ∼ 3000 s. Each dot corresponds to one recording, and colors correspond to (a).

**Figure S4:**
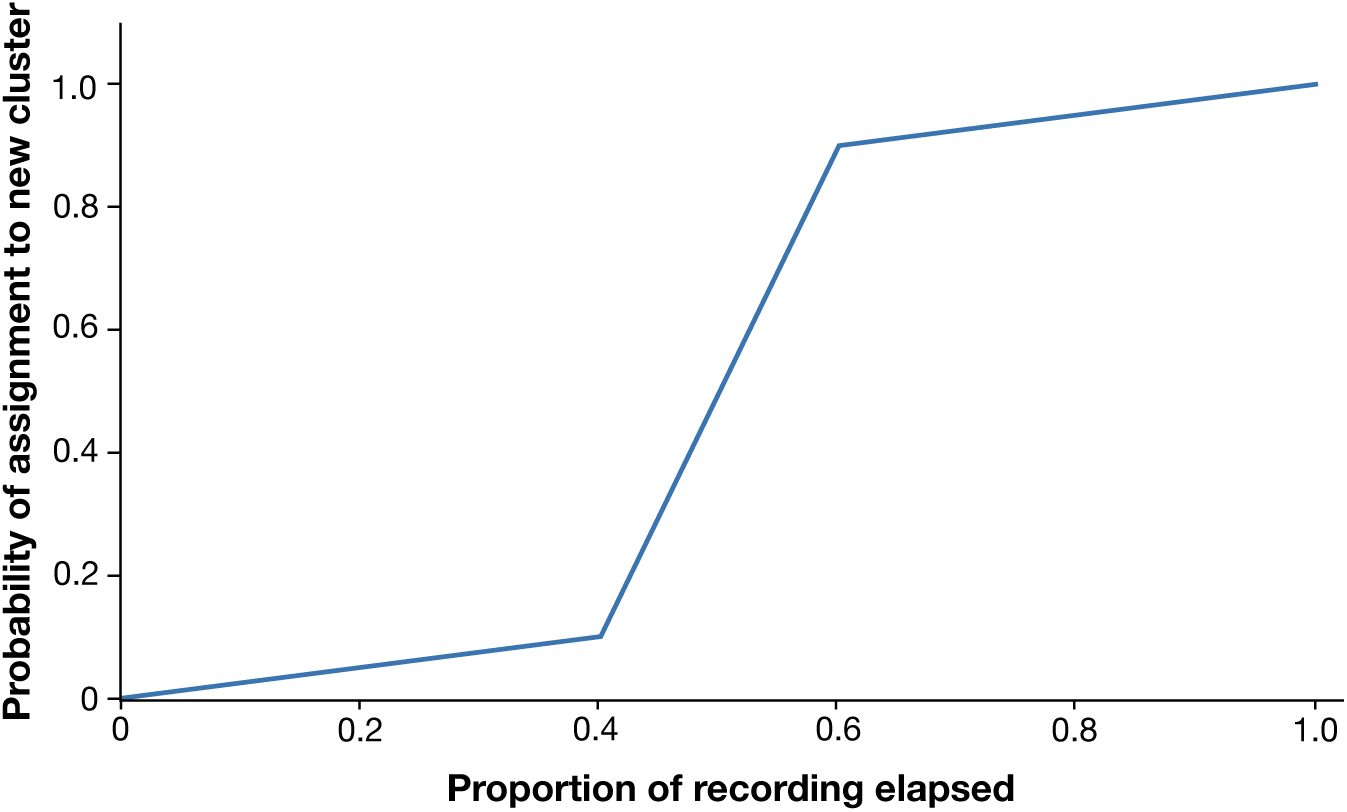
Split probability function for artifical drift splits. Spikes in the first 40% of the recording have a 0-10% chance of being assigned to the new unit, spikes in the middle 20% have a 10-90% chance, and spikes in the last 40% have a 90-100% chance. This more realistically models drift-related errors, which are a gradual process rather than a sudden change in spike assignment.

**Figure S5:**
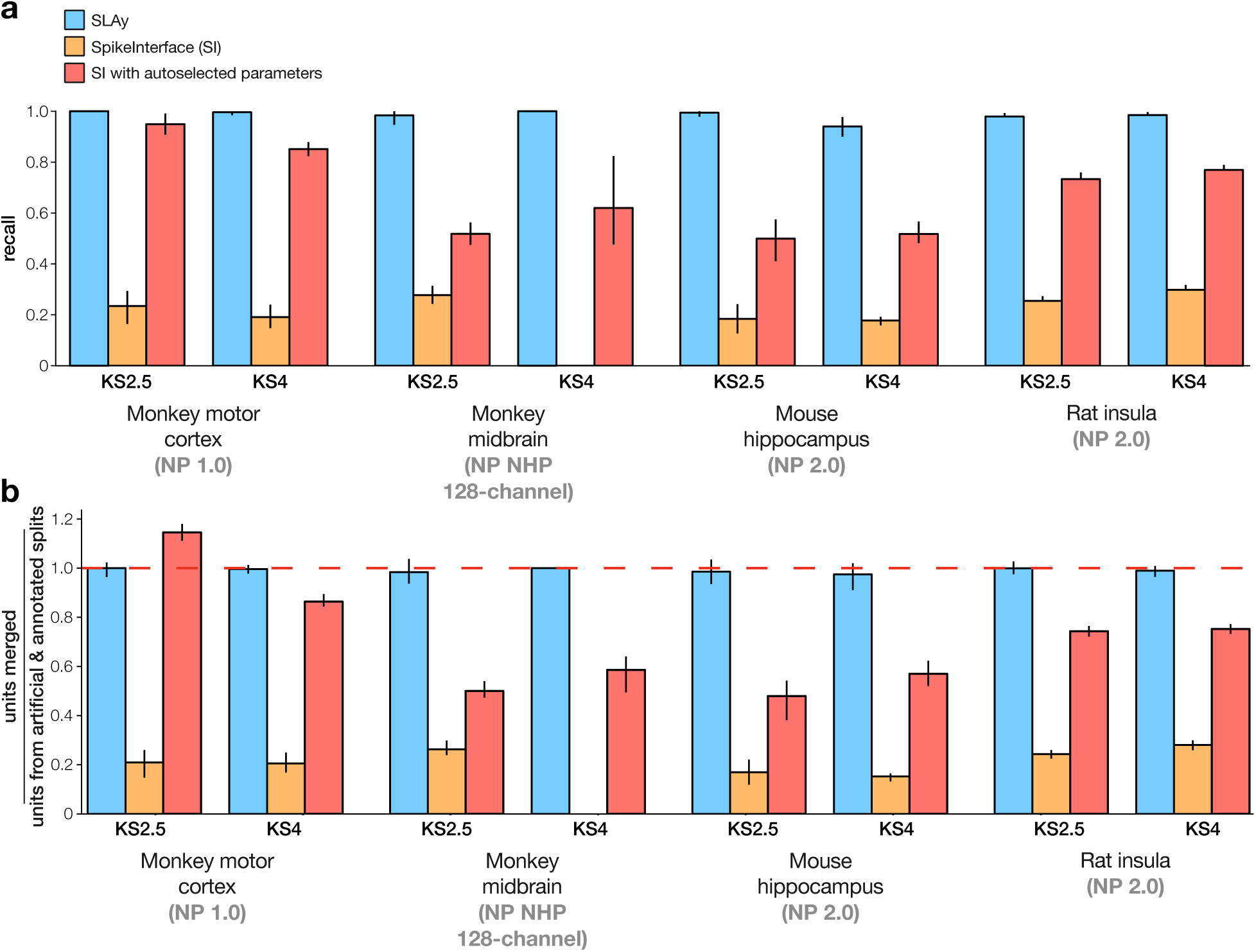
Per-dataset performance on simulated oversplitting. **(a)** Recall on simulated oversplitting for SLAy (blue), similarity-correlograms (SC, orange), and SC with autoselected parameters (SC-auto, red), broken down by recording and Kilosort version. Error bars show the 95% confidence interval across five independent draws of simulated oversplitting. SLAy achieves near-perfect recall across all datasets, while SC and SC-auto show substantially lower recall. **(b)** Ratio of units merged by each algorithm to the total number of split units (artificial and curator-identified). A ratio of 1 (dashed red line) indicates merging exactly the expected number of units; ratios above 1 indicate overmerging. SLAy consistently stays near 1 across all recordings, while SC-auto exceeds 1 in the KS2.5 monkey motor cortex dataset. SC with default parameters is overly conservative, consistently merging far fewer than the expected number of merges. Bars show means across five draws; error bars show 95% confidence interval.

**Figure S6:**
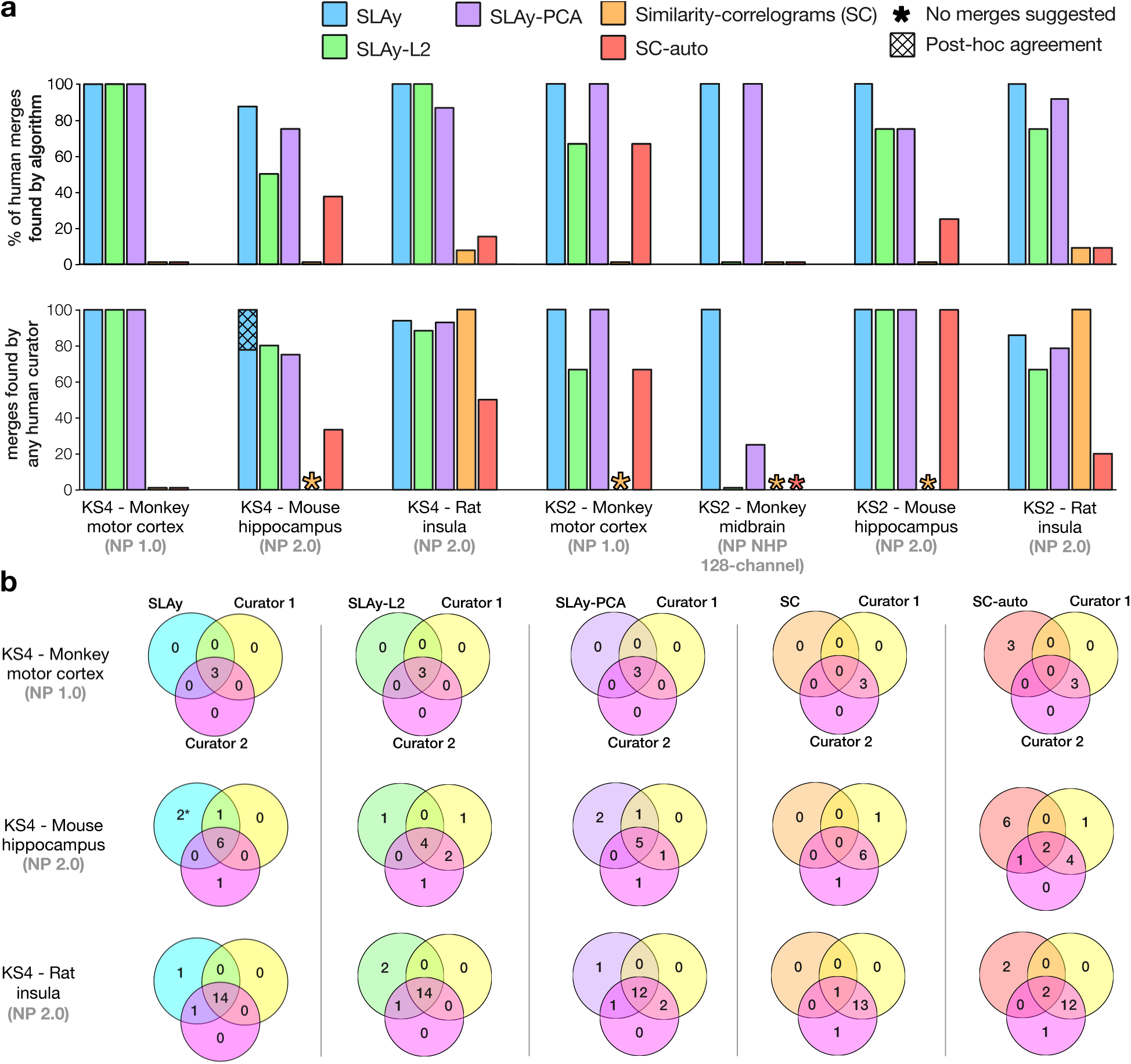
Per-dataset comparison of automated merging algorithms to human curators. **(a)** Recall (top; percent of human curator merges recovered by each algorithm) and precision (bottom; percent of algorithm merges agreed with by a human curator) for each merging algorithm on each dataset. Improved precision for merges validated by a human curator during post-hoc blinded examination is shown as a hatched bar. indicate no merges were suggested by the algorithm for that dataset. **(b)** Venn diagrams showing the overlap in merges between each algorithm and two human curators for each KS4 recording.

**Figure S7:**
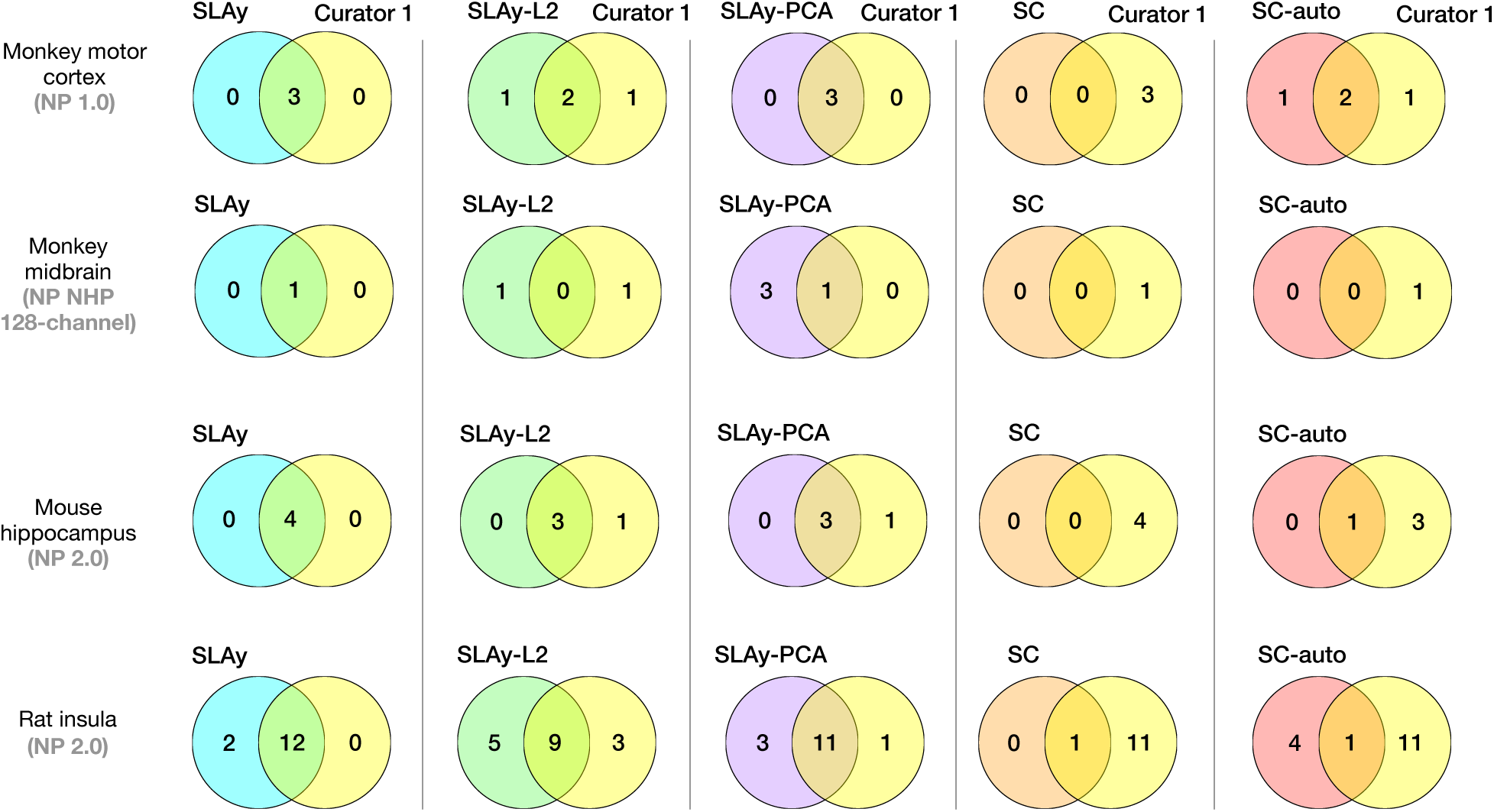
Comparison of automated merging algorithms to human curators on Kilosort 2.5. Venn diagrams showing the overlap in merges between each algorithm and the single human curator for each KS2.5 recording.

**Figure S8:**
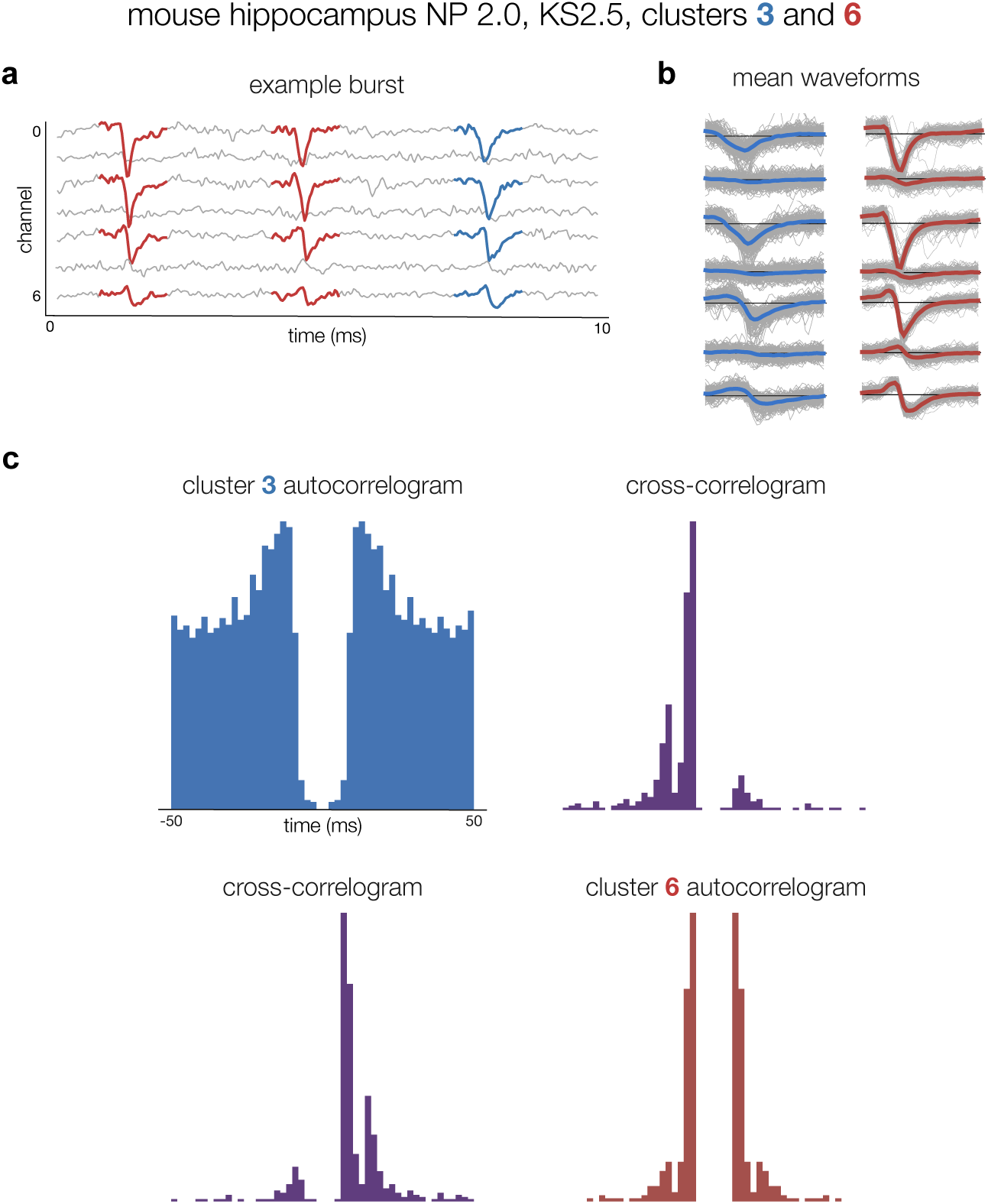
SLAy accounts for biophysical waveform changes. SLAy merges two units in the mouse hippocampus recording that were oversplit by Kilosort 2.5 due to bursting-related waveform changes. **(a)** A salient burst in the recording was split into two different units. The first two spikes from the recording were assigned by Kilosort to unit 6 (red), and the last spike in the burst had a wider waveform shape, causing it to be assigned to unit 3 instead (blue). **(b)** The mean waveforms for the two units show significant differences (colored traces), though the distributions of individual spikes (thin grey traces) overlap between units. **(c)** The cross-correlogram has a highly peaked and asymmetric shape, characteristic of an oversplit bursting neuron.

**Figure S9:**
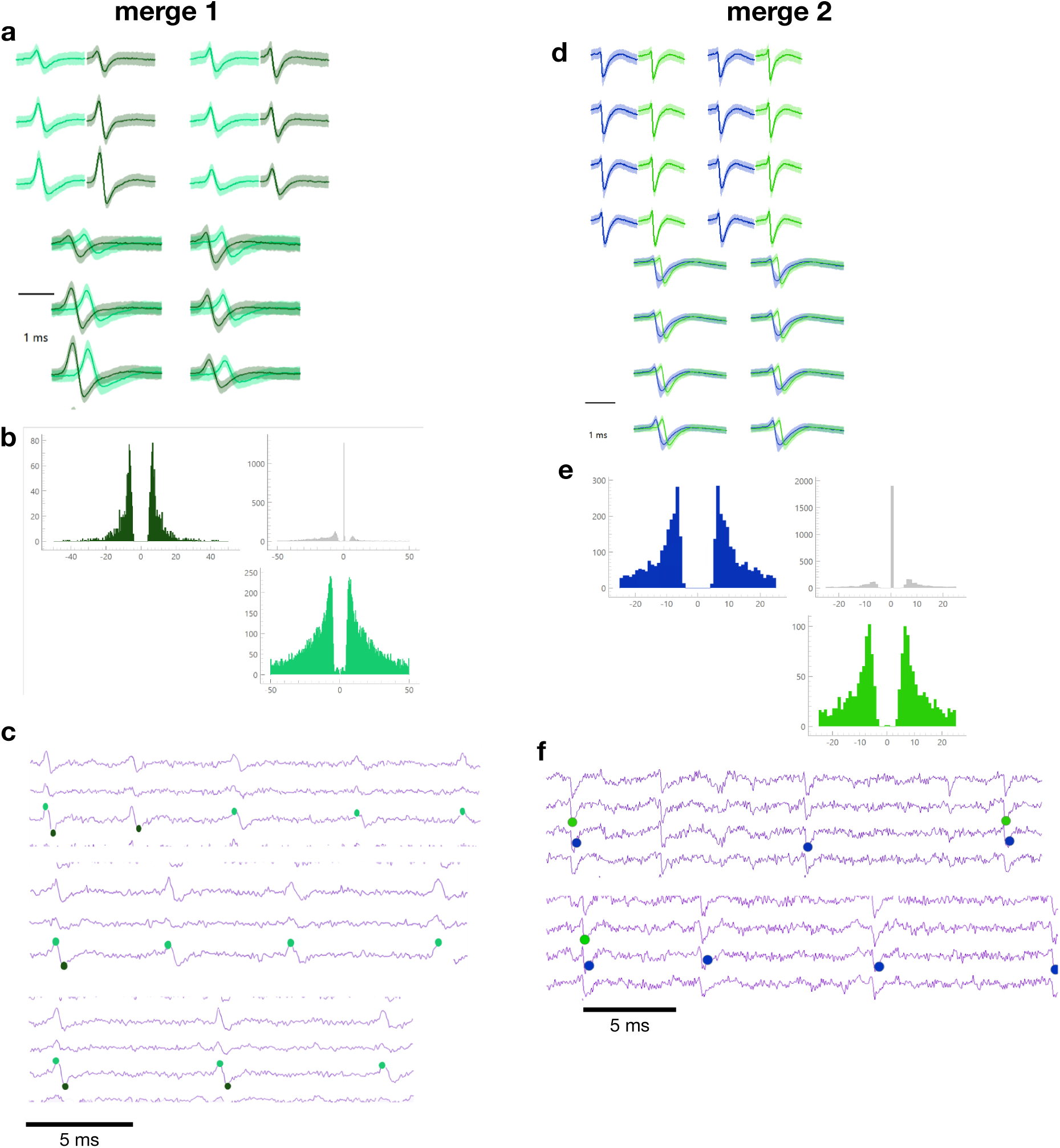
SLAy identifies oversplit units that human curators initially missed. Two example unit pairs that SLAy flagged for merging. Human curators missed these initially, as the pairs did not pass the curation GUI’s initial L1 similarity screening step, but a blinded human curator agreed with the merges post-hoc. **(a)** Mean waveforms for the units in merge 1, plotted separately (top) and overlapped (bottom). Shading indicates one standard deviation across individual spike waveforms. **(b)** Auto- and cross-correlograms for the units in merge 1, showing a large central peak, consistent with a co-detected spikes error, and asymmetry, consistent with an oversplit burst neuron. **(c)** Example voltage traces with colored dots showing the detected spike times for each unit. Within a single burst, some spikes are detected with both units’ templates, some with only one, and some with only the other. This indicates that the units each capture unique information about a ground-truth bursting neuron, and that merging them together is vital for accurate analysis of the neuron’s spiking. **(d-f)** Same as (a-c) for another merge pair found by SLAy but missed by human curators, with similar characteristics to the first.

**Figure S10:**
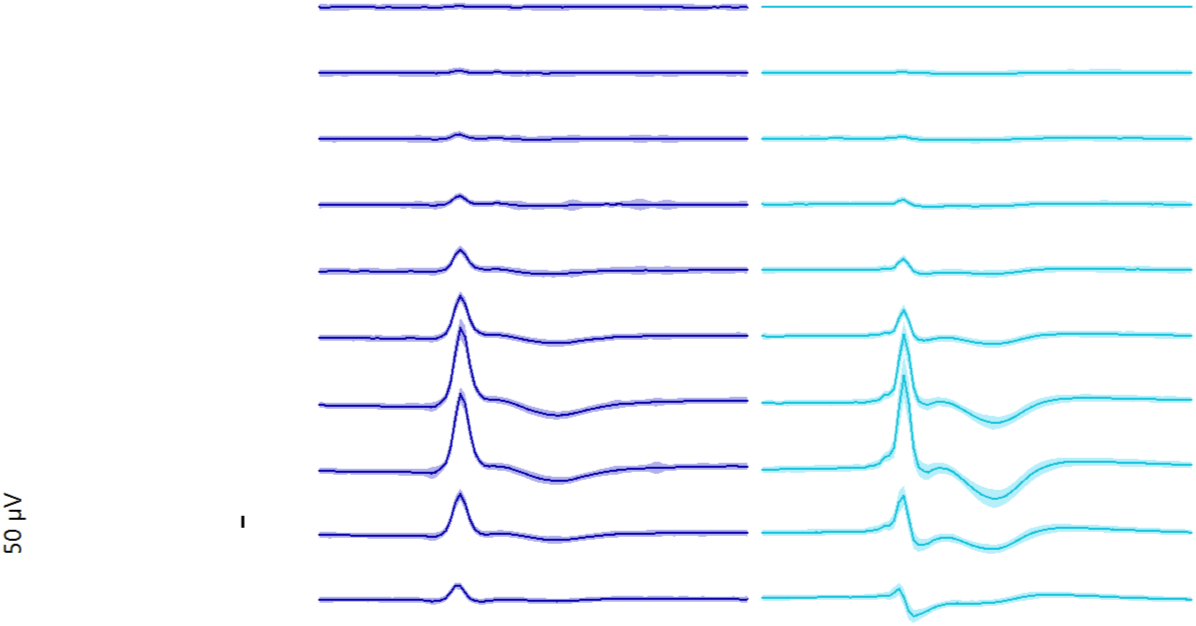
Example false-positive merge suggested by SLAy. Mean waveforms for a pair of large-amplitude units (blue and cyan) across multiple electrode channels (50 µV scale bar) that SLAy suggested merging but a human curator disagreed with upon blinded re-evaluation. The two units have similar overall waveform shapes but differ subtly in the repolarization phase, most clearly visible on the highest-amplitude channels. Discrepancies appear to be exceed the normal variation for each unit (shaded region; standard deviation). The autoencoder likely did not learn sufficiently distinct latent representations for the two units, resulting in an erroneously high waveform similarity score.

**Figure S11:**
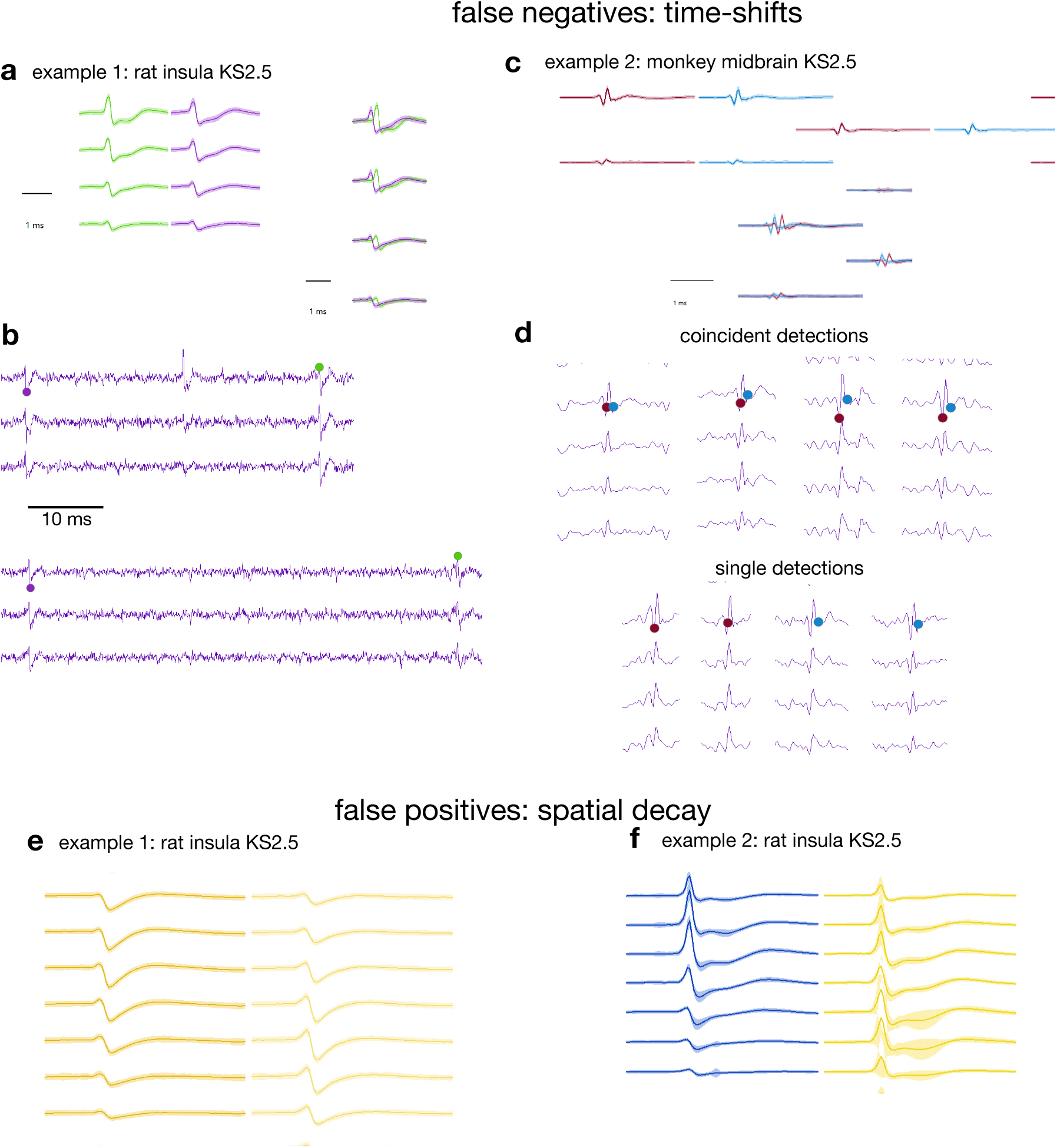
SLAy-L2 makes stereotyped errors due to limitations of L2 waveform similarity. (a-d) False negative examples in which SLAy-L2 failed to merge temporally shifted units. **(a)** Multi-channel mean waveforms across channels for two units from the rat insula KS2.5 recording that are shifted by a few samples relative to each other; L2 distance squares and sums sample-wise differences, inflating the apparent dissimilarity for shifted waveforms. **(b)** Raw voltage traces with the spike times of each unit marked (colored dots). Spikes detected with one unit’s template vs. the other do not show a clear visual difference, suggesting that both units represent the same underlying neuron. **(c)** Mean waveforms for a second temporally shifted pair from the monkey midbrain KS2.5 recording. **(d)** same as (b) for the monkey midbrain example, again showing no discernible difference for spikes detected with both or only one unit’s template. **(e–f)** False-positive examples in which SLAy-L2 erroneously merged units with differing spatial decay profiles. Although the two units in each pair have similar waveforms on the highest-amplitude channels, their waveforms decay differently across the probe. L2 distance is dominated by high-amplitude channels and fails to penalize spatial differences; SLAy’s autoencoder, which must reconstruct the multichannel waveform, better distinguishes such pairs.

**Figure S12:**
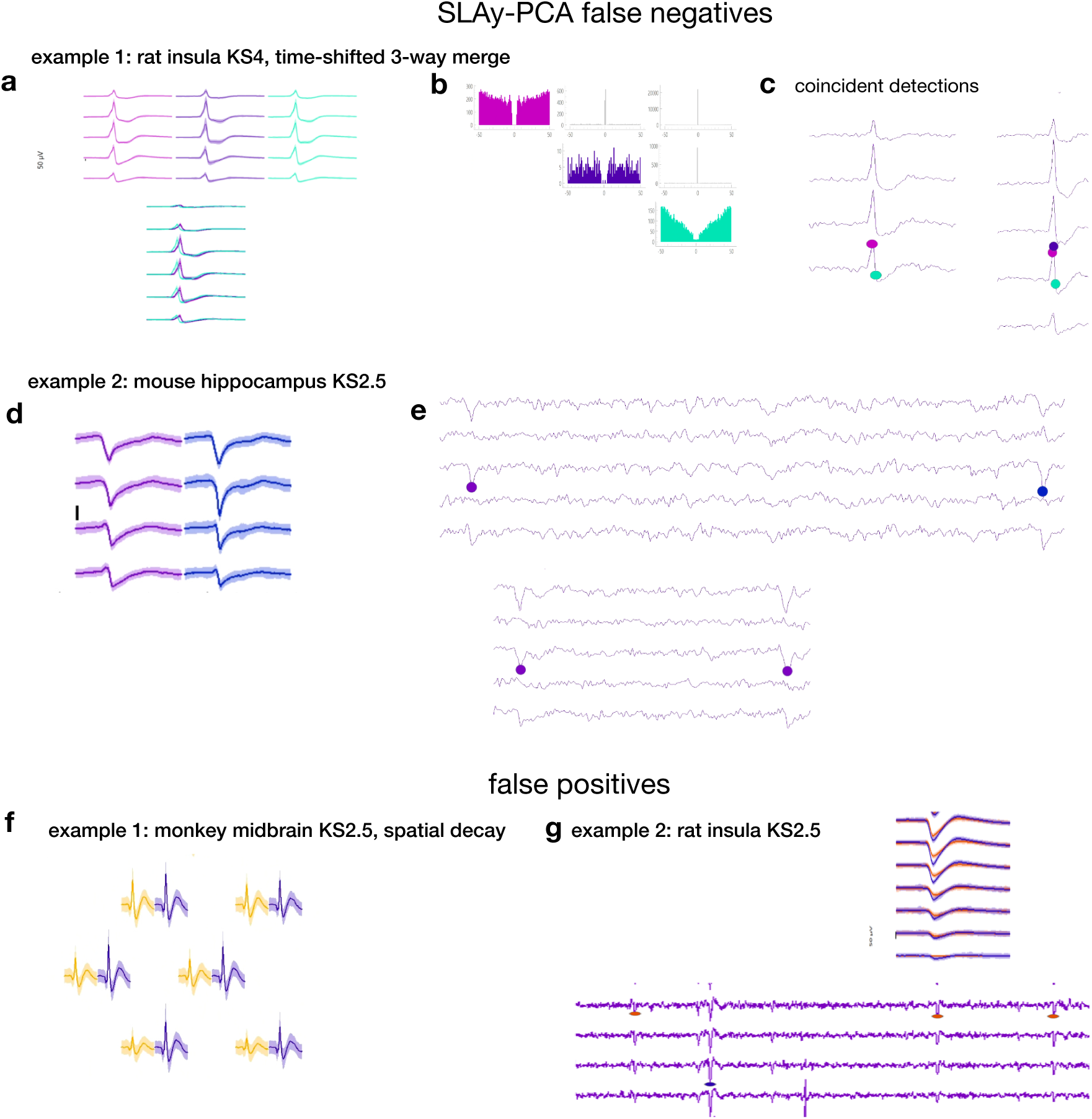
PCA features result in false positives and false negatives. (a-c) A three-way merge from the rat insula KS4 recording. Two human curators, SLAy with autoencoder similarity, and SLAy-L2 independently identified this merge. SLAy-PCA merged only the pink and purple units, but not the turquoise unit, presumably due to the slight timeshift between this unit and the other two. **(a)** Mean waveforms for the units involved in the merge, plotted separately (top) and overlapping (bottom). **(b)** Auto- and cross-correlograms for the three units. CCGs show a large central spike consistent with frequent co-detection of a single spike with multiple unit templates. For these units, a very high proportion (*>* 90%) were co-detected. **(c)** Multi-channel traces from reprsentative example spikes from these units. Colored circles denote a detection of a spike with the corresponding unit’s template. Close circles of different colors in the examples show co-detection of individual spikes with multiple units’ templates. **(d-e)** A merge from the mouse hippocampus KS2.5 recording that was missed by SLAy-PCA but found by SLAy and an independent human curator. **(d)** Multi-channels mean waveforms, showing highly similar waveform shapes and overlapping amplitude distributions (shaded). **(e)** Representative example traces showing spikes detected with each unit’s template. Spikes detected with different templates are not visually different. **(f)** Multi-channel mean waveforms of a false positive example from the monkey midbrain KS2.5 dataset, in which SLAy-PCA erroneously merged units with different spatial decay across sites. The yellow unit’s amplitude decays considerably from the peak channel (top left) to the lowest-amplitude channel (bottom right), while the purple unit’s amplitude decays only slightly. **(g)** A false positive example from the rat insula KS2.5 dataset, in which SLAy-PCA merged two units with tight, separable amplitude distributions (top) and clear visual differences between detected spike traces (bottom). The sequence of spike amplitudes in the bottom trace is not consistent with biophysical changes in a single neuron’s waveform (e.g. monotonic decrease due to bursting)

**Figure S13:**
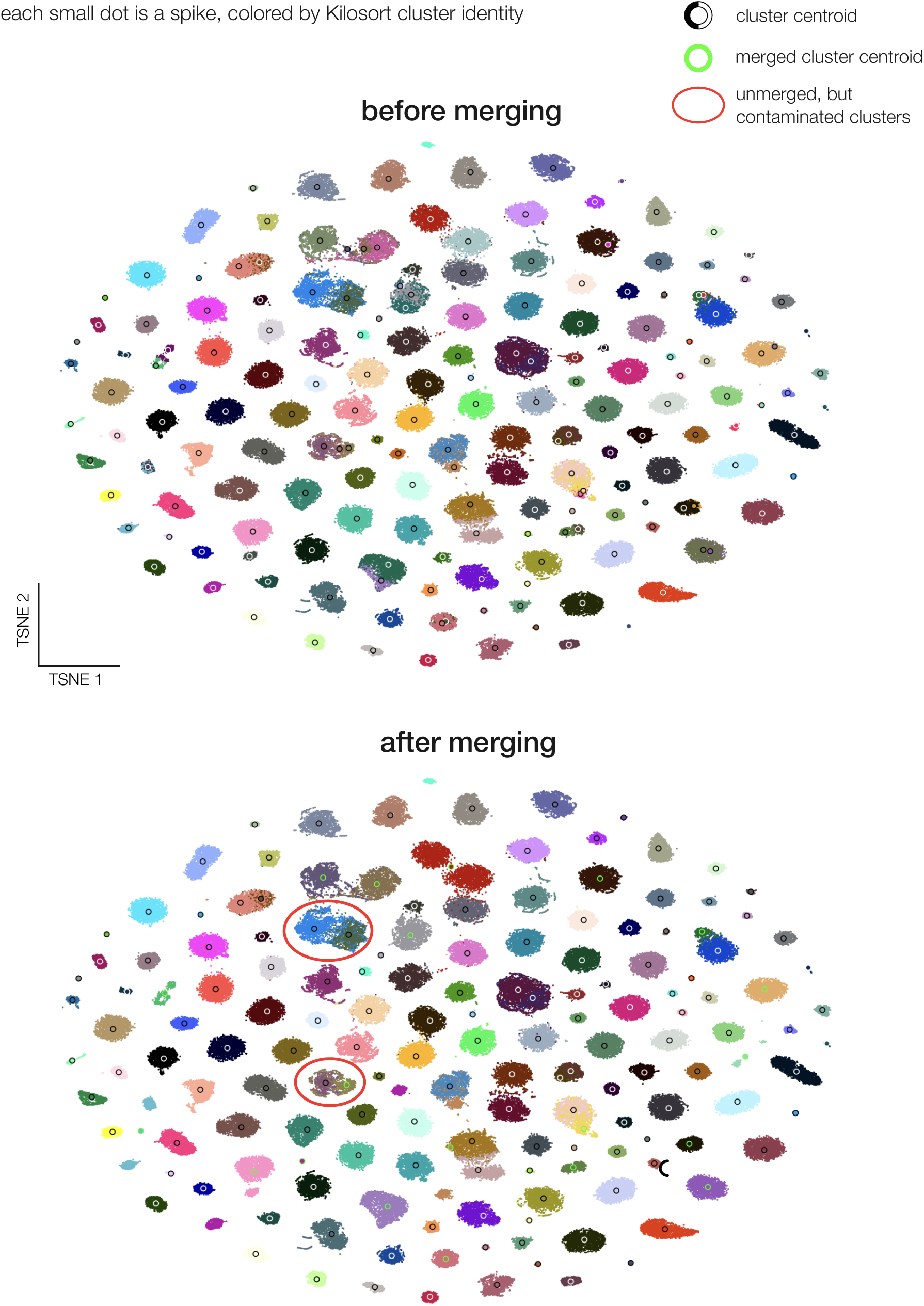
SLAy merges units that are close together in waveform space. TSNE plots of autoencoder latent space colored by unit ID, befor_4_e_3_(top) and after (bottom) merging. The green unit centroids show that only units that are close together in latent space were merged together. Red circles show units that are close together in waveform space, but were not merged together, presumably because of refractory period violations.

**Figure S14:**
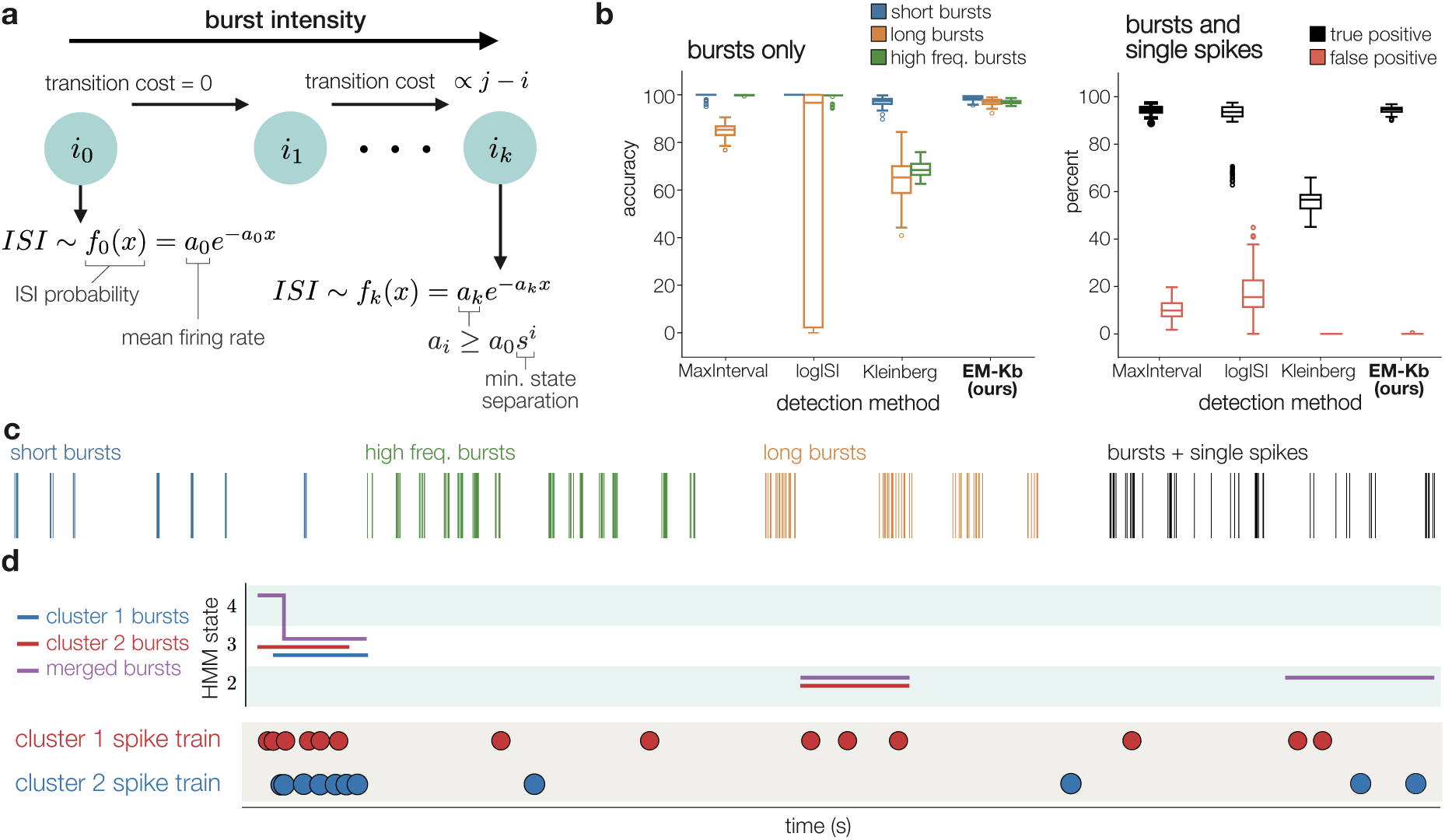
Robust burst detection. **(a)** Our burst detection algorithm relies on a Hidden Markov model where each state corresponds to a different firing rate, and adjacent states have firing rates separated by at least a scaling factor of *s*. Over the course of a spike train, neurons transition to higher states when bursting and lower states when firing normally. We learn both the state sequence and per-state firing rates using an expectation-maximization algorithm. **(b)** Our burst detection algorithm outperforms three existing methods on simulated spike trains with a variety of burst statistics. **(c)** Example simulated spike trains with different bursting properties. **(d)** Applying our burst detection algorithm before and after using SLAy shows that correcting oversplitting errors is necessary for accurate downstream analysis. After merging, our bursting algorithm detects additional bursts and estimates a higher intensity for the already detected bursts.

**Figure S15:**
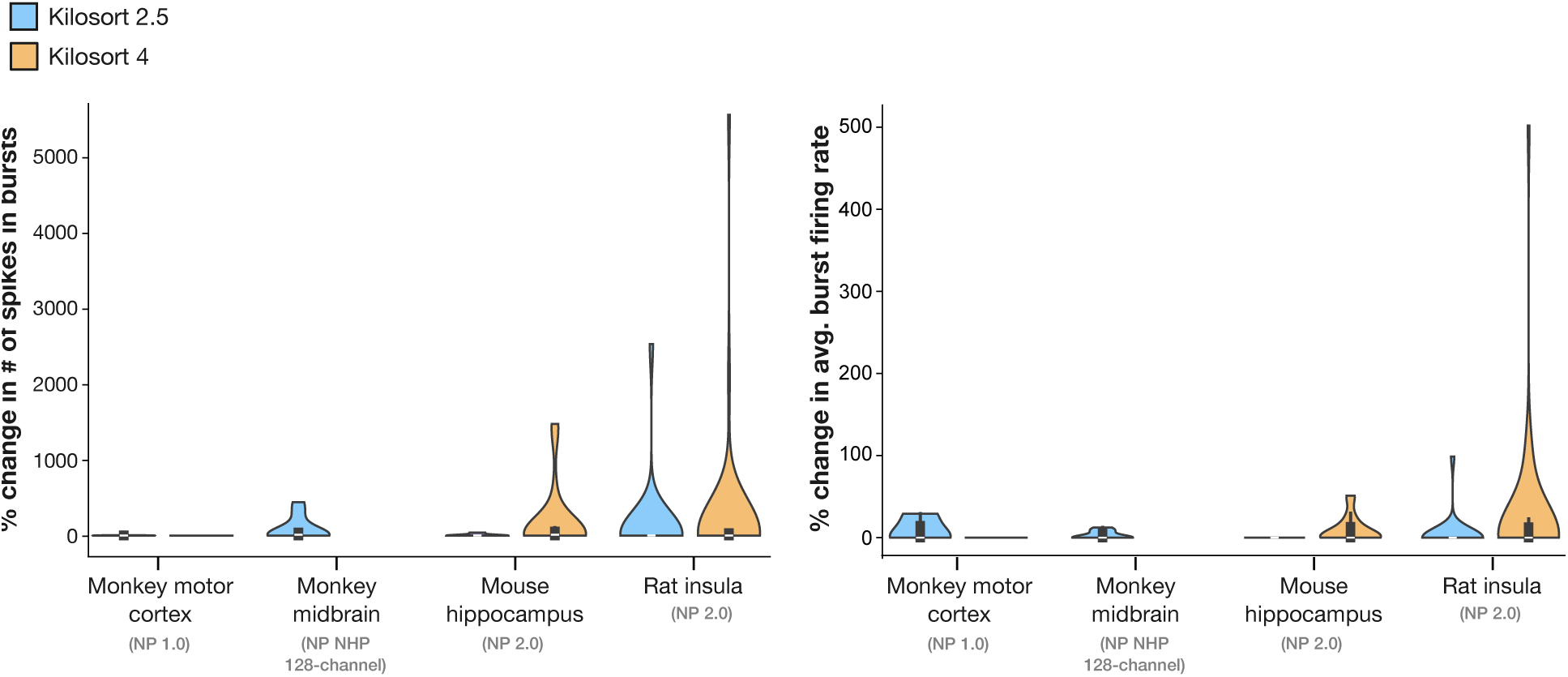
Per-dataset effect of SLAy merging on burst detection metrics. Violin plots of the percent change in the number of spikes in bursts (left) and average burst firing rate (right) following SLAy merging, broken down by recording and Kilosort version (blue: KS2.5; orange: KS4). Each data point corresponds to one merged unit. The largest increases in both metrics are observed in the mouse hippocampus and rat insula datasets, consistent with high rates of bursting activity in these regions. The monkey motor cortex shows minimal change, corresponding to the small number of merges identified in that dataset.

## Notes

https://github.com/saikoukunt/SLAy

### Competing Interest Statement

The authors have declared no competing interest.

### Summary of Updates

This revision includes additional analyses and comparisons to other algorithms, as well as a automated parameter fitting method to ease usability.

